# Remote Activation of Wnt Signaling and Cell Proliferation by E-cadherin Magnetomechanical Stimulation

**DOI:** 10.64898/2026.01.08.698380

**Authors:** Christian Castro-Hinojosa, Pablo Martínez-Vicente, Susel Del Sol-Fernández, Pilar Gomollón-Zueco, Yilian Fernández-Afonso, Lucía G. Recaredo, Raluca M. Fratila, María Moros

## Abstract

The ability to remotely and precisely manipulate intracellular signaling pathways is a powerful tool for both fundamental biological research and therapeutic applications. Among these pathways, the Wnt/β-catenin signaling cascade plays a central role in regulating cell proliferation, differentiation, and tissue regeneration. However, current methods for activating this pathway such as pharmacological agents lack spatiotemporal control and may induce severe off-target effects. In this study, we introduce a pioneering magnetogenetic toolkit to modulate the Wnt/β-catenin pathway through magnetomechanical stimulation of E-cadherin, a key cell adhesion molecule intimately linked to β-catenin dynamics.

Engineered magnetic nanoparticles (MNPs) functionalized with the extracellular domain of E-cadherin (MNPs@E/EC15) are used to selectively bind cellular E-cadherins. By applying a weak intensity and low-gradient magnetic field using a custom-designed magnetic stimulator, localized mechanical forces sufficient to trigger E-cadherin-mediated mechanotransduction are produced. This stimulation leads to β-catenin release from the membrane, nuclear translocation, and activation of Wnt target gene expression, as confirmed by transcriptomic profiling and a Wnt-responsive luciferase reporter assay. These molecular changes are also translated into functional outcomes, including enhanced cell proliferation and accelerated wound closure.

This work establishes an innovative non-invasive tool for probing E-cadherin mechanobiology and remotely modulating Wnt/β-catenin signaling with high spatiotemporal resolution. Unlike other tools to probe mechanotransduction, this approach enables the simultaneous modulation of many cells with precise control, using low intensity magnetic field that could be potentially translated into *in vivo* designs. Our findings open promising avenues for studying mechanotransduction and developing targeted regenerative therapies based on mechanical stimulation.

---

The possibility of externally manipulating intracellular signaling pathways with spatio-temporal control is among the most promising tools for understanding basic biology and for developing clinical applications.^1,2^ This control is often achieved through the activation of cell membrane receptors, which serve as key starting point for intracellular signaling cascades that ultimately regulate gene expression and determine specific cell responses.^3–5^ Different physical tools involving the use of light, electricity, ultrasound, or magnetic fields have been developed to deliver non-invasive stimuli to manipulate intracellular responses.^6–8^ Notably, several signaling pathways can also be activated through the mechanical stimulation of specific mechanosensors, including mechanosensitive ion channels^9^ and other membrane proteins.^10^ Mechanical stimulation is increasingly recognized as a crucial mechanism by which cells interpret the environment, finally impacting numerous cellular processes.^5^ In particular, the use of magnetic actuators alongside magnets to remotely modulate cellular activity or gene expression, which is coined as magnetogenetics,^6,11^ has opened new avenues for the controlled manipulation of intracellular pathways, showing high potential for *in vivo* applications.^3^

Among these signaling cascades, the Wnt pathway stands out as an attractive target for precise on-demand regulation for tissue engineering. This evolutionarily conserved cell-cell communication system governs essential cellular functions such as differentiation, proliferation, embryonic development, and tissue homeostasis.^12,13^ Wnt signaling is divided into canonical and noncanonical pathways. The canonical pathway involves the ligand recognition by Frizzled receptors leading to the subsequent nuclear translocation of β-catenin, a key cytoplasmatic effector that allows the transcription of target genes *via* T-cell factor/lymphoid enhancer-binding factor (TCF/LEF) transcription cofactors.^14^ By contrast, non-canonical pathways are independent of β-catenin, and include the Wnt/Ca^2+^ and the planar cell polarity pathways.^15^

Under physiological conditions, canonical Wnt/β-catenin signaling is meticulously regulated and plays a beneficial role in processes such as wound healing and tissue regeneration.^12,13,16^ However, an uncontrolled activation can lead to unrestricted proliferation and the onset of cancer.^17,18^ Achieving precise control over Wnt/β-catenin signaling activation remains a major challenge, particularly for regenerative medicine applications where targeted tissue regeneration is only desired in specific areas. Current approaches, such as chemical modulation using compounds like BIO (6-bromoindirubin-3’-oxima; GSK-3 inhibitor) or recombinant Wnt3a protein lack spatiotemporal control, potentially leading to off-target effects.^18–21^ On the other side, the use of optogenetics to control this pathway requires the genetic engineering of the cells to express light-sensitive constructs.^22^ Therefore, new approaches that enable remote and localized control of the Wnt/β-catenin pathway are needed.

In this work we propose a completely unprecedented strategy to manipulate the Wnt/β-catenin pathway in a specific manner, taking advantage of its close functional linkage with cadherins.^23^ E-cadherin constitutes the principal adhesion protein of the cell-to-cell contact machinery, and its mechanotransduction is critical to mediate collective epithelial remodeling that takes place during tissue repair.^24^ Aside from its potential as specific cellular targets, E-cadherins bind in its cytoplasmatic tail to β-catenin, which is the central player of the canonical Wnt pathway. While initially E-cadherin was considered a negative regulator of canonical Wnt pathway (as it prevents β-catenin nuclear translocation by maintaining it at the membrane),^25,26^ recent studies have demonstrated that E-cadherin can modulate β-catenin dynamics in a more complex manner, playing a crucial role in Wnt/β-catenin signaling regulation. As such, stretching of E-cadherin with a force of 6 pN has been shown to induce a conformational change in β-catenin, leading to its phosphorylation and irreversible dissociation from E-cadherin.^27^ Similarly, sustained biaxial stretching resulted in the translocation of β-catenin to the nucleus in an E-cadherin-dependent manner, promoting cell cycle entry.^28,29^ This provides direct evidence of the link between E-cadherin mechanical stimulation, β-catenin signaling, and cell proliferation.

Given this interplay, the mechanical stimulation of E-cadherin emerges as a promising strategy for modulating the Wnt/β-catenin pathway and its downstream transcriptional responses associated with cell proliferation.^24,30^ In this context, various technologies such as atomic force microscopy (AFM) and optical or magnetic tweezers have been successfully employed to study the effects of tensile forces on cadherins.^31–33^ However, these tools present significant limitations: they cannot be applied remotely, suffer from limited penetration for *in vivo* applications and/or typically allow stimulation of only one or few cells at a time. Conversely, macroscopic techniques such as biaxial stretching or shear stress can stimulate simultaneously a larger number of cells but face other hurdles. For instance, monolayer stretching is an interesting approach as it mimics the physiological mechanical cues experienced by the cells during processes like tissue remodeling, offering a more accurate representation of the mechanotransduction signaling. However, it applies indirect and non-uniform forces across entire cell sheets, leading to global mechanical stimuli that can potentially affect multiple mechanotransduction pathways.

To overcome these challenges, magnetogenetics, that uses magnetic fields in combination with magnetic actuators, has become an invaluable tool, enabling the stimulation of mechanoreceptors with exceptional spatiotemporal control even *in vivo*.^6^ Among the different magnetic actuators, MNPs offer higher spatial resolution than larger microparticles;^11^ however, their smaller size limits the magnitude of mechanical forces they can exert, which makes necessary a careful optimization of both the MNP properties and the magnetic field applicator to reach the force threshold required to activate the mechanoreceptor. While cadherin mechanotransduction has been extensively demonstrated using magnetic particles,^34–37^ to the best of our knowledge, it has not been studied to selectively activate the Wnt/β-catenin pathway, nor for cellular proliferation.

In this study, we developed a novel nanoplatform to activate with high precision the Wnt/β-catenin pathway through the mechanical stimulation of E-cadherins, isolating the effects promoted specifically by E-cadherin stimulation from those produced by other mechanotransduction pathways. To this end, MNPs were functionalized with the extracellular domain of E-cadherin (E/EC15) to create a nanoactuator capable of selectively recognize cellular E-cadherins. By applying a low intensity magnetic field using a customized magnetic field applicator, these bioconjugates exerted enough mechanical forces on the cellular E-cadherin to modulate β-catenin dynamics in epithelial cells. This led to the activation of the Wnt/β-catenin signaling pathway and enhanced cellular proliferation (Figure 1). We performed a comprehensive set of experiments, including RNA-seq analysis, protein expression assays, and functional proliferation studies to elucidate the mechanistic link between E-cadherin mechanotransduction and Wnt/β-catenin signaling. Noteworthy, we demonstrate for the first time that torque forces generated by MNPs can activate this pathway through E-cadherin stimulation, introducing a novel and precise approach for remotely modulating cellular responses with high spatiotemporal resolution. The use of low magnetic fields can be easily translated to *in vivo* settings, opening new avenues in mechanotransduction and regenerative medicine, where controlled cell stimulation is essential.

**Figure 1.**
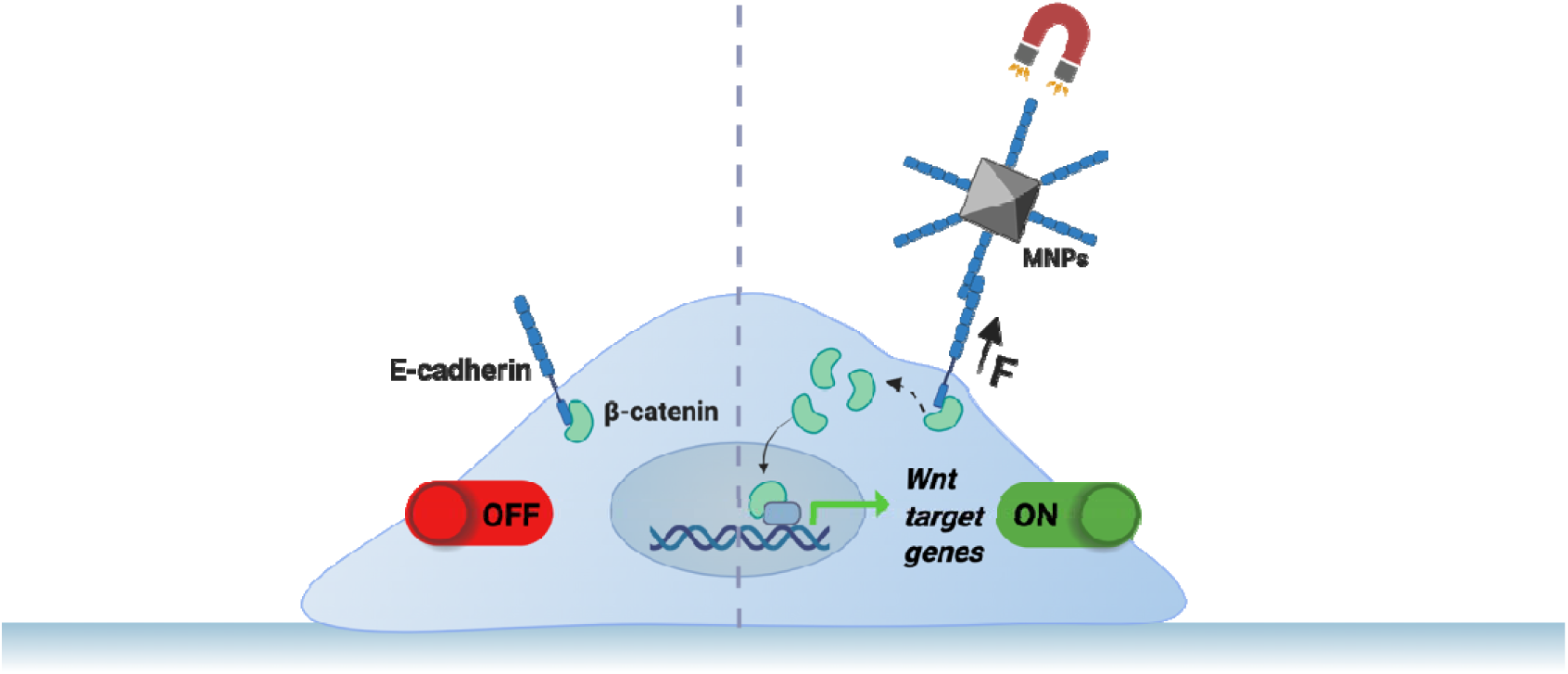
Schematic overview of the general concept of Wnt/β-catenin pathway activation *via* E-cadherin mechanostimulation. In the absence of force, β-catenin remains anchored to the cytoplasmatic tail of E-cadherin at the cell membrane. Conversely, upon targeting cellular E-cadherin with MNPs functionalized with its extracellular domain and applying a weak magnetic field, torque forces are generated on the MNPs. These forces promote the dissociation of β-catenin from the E-cadherin complex into the cytoplasm and its subsequent translocation into the nucleus, where it activates Wnt target gene transcription. Image created in https://BioRender.com.

## Results & Discussion

### Toolkit generation

#### Production of E-cadherin Fragments E/EC15

Cellular cadherins are composed of five extracellular domains (E/EC15) containing four N-glycosylation sites (asparagine residues 554, 566, 618, and 633) that mediate calcium-dependent homophilic interactions. While particles functionalized with antibodies are frequently used to target cellular cadherins with high affinity, the biological response they elicit may differ from those mediated by native cadherin-cadherin interactions.^34,38^ For instance, only beads coated with E-cadherin fragments were able to modulate cell stiffness under mechanical stimulation; in contrast, microparticles decorated with antibodies failed to induce this response.^39^ This result suggests that effective cadherin mechanical stimulation needs to be performed using ligands that closely mimic their native homophilic interactions.

Our group has recently described the possibility to selectively target cellular cadherins using MNPs modified with cadherin fragments comprising only the two outermost E-cadherin domains (E/EC12).^40^ However, these fragments, produced in *Escherichia coli,* present several drawbacks: i) bacterial expression leads to the production of cadherin fragments with a methionine residue as the first translated amino acid, which could partially affect the interaction affinity of these fragments with the cellular cadherins;^41,42^ ii) bacterial expression lacks post-translational modifications, potentially compromising protein stability and adhesion properties,^43^ iii) it has been demonstrated that the immobilization of the full extracellular domain of N-cadherin (N/EC15) on hydrogels promotes higher secretion of key proteins involved in mesenchymal stem cell differentiation than shorter fragments (N/EC12), highlighting the functional importance of larger sequences.^39^ Moreover, regarding MNP bioconjugation, longer E/EC15 fragments are expected to project the adhesive EC1 domain further from the MNP surface and the PEG grafting, thereby reducing any steric hindrance and improving accessibility to cellular cadherins compared to E/EC12 fragments. Thus, to better mimic native cadherin-cadherin interactions, the full E-cadherin extracellular domain E/EC15 was produced in mammalian cells, which support the expression of larger proteins while preserving essential post-translational modifications, such as pro-peptide cleavage and both O and N-glycosylations.

E/EC15 fragments containing a C-terminal 6-histidine tag^44^ were purified using immobilized metal affinity chromatography (IMAC) for His-tagged proteins.^45^ Sodium dodecyl-sulfate polyacrylamide gel electrophoresis (SDS-PAGE) analysis revealed a single band at ∼65 kDa in the first recovered fraction (Figure 2a), corresponding to the theoretical molecular weight of E/EC15 (61 kDa). To identify the obtained band, liquid chromatography-electrospray ionization-mass spectrometry (LC-ESI-MS) analysis was performed following a trypsin digestion, allowing peptide comparison with public protein sequence databases (Swissprot, NCBI).^46,47^ The highest protein score was attributed to E-cadherin (score: 1185) (Figure S1, Supporting Information).^48^ This result confirmed the successful production and purification of mature E/EC15 fragments.

**Figure 2.**
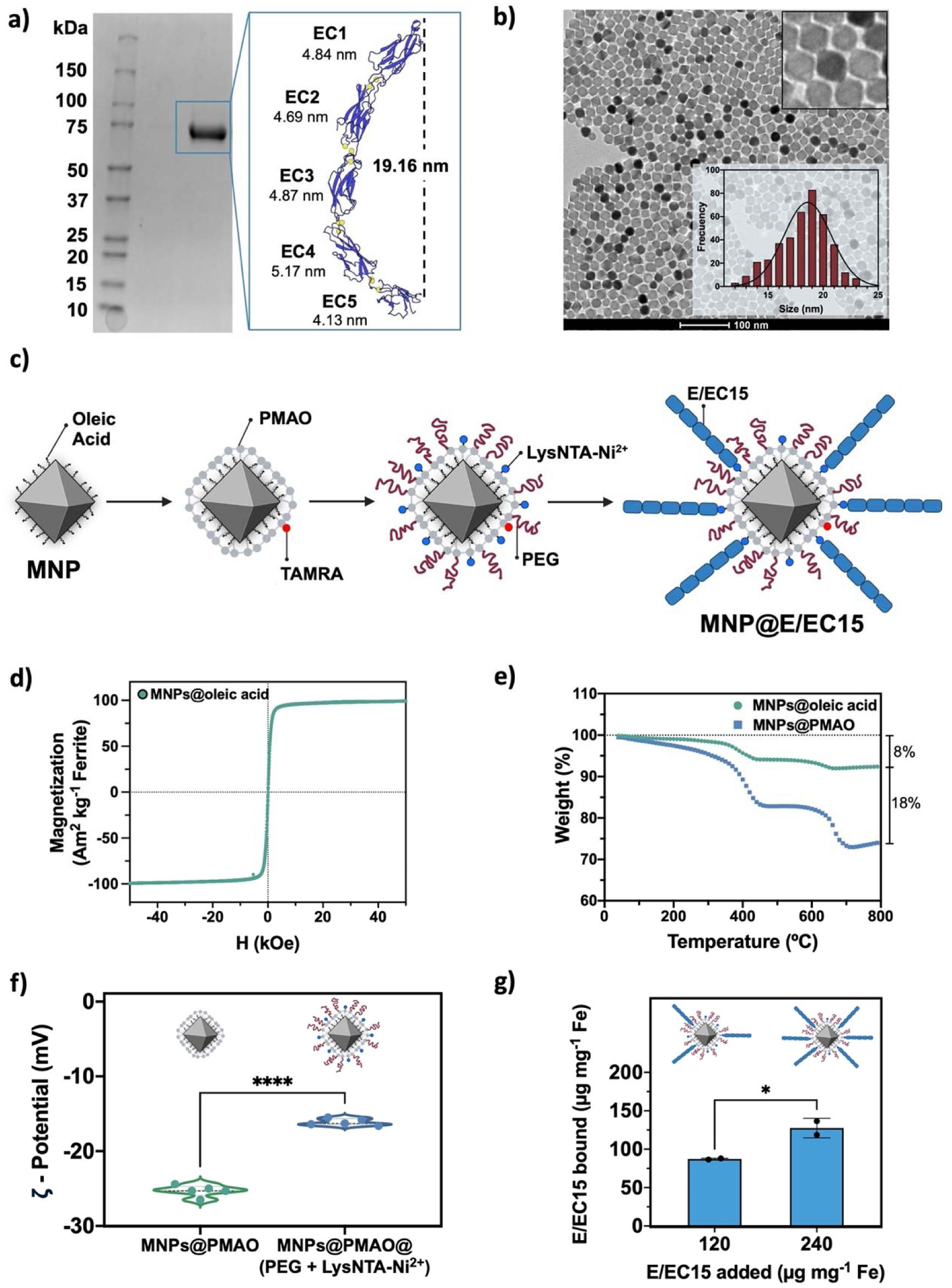
Synthesis and characterization of MNPs@E/EC15. a) SDS-PAGE of the E/EC15 fragments. The band at ∼65 kDa corresponds to the theoretical molecular weight of E/EC15. Schematic representation of the fragment, conformed by five extracellular domains EC1-EC5 (in blue) and calcium ions (in yellow) (approximate sizes were determined from PDB database using the PyMOL software). The protein scheme was created in Protein Imager.^52^ b) TEM image showing the synthesized octahedral MNPs and their edge size distribution in hexane fitted with a Gaussian function (350 particles measured); scale bar: 100 nm. c) Scheme of the functionalization steps to obtain MNPs@E/EC15. Hydrophobic MNPs (MNPs@oleic acid) were coated with PMAO (previously modified with TAMRA), then functionalized in a single step with PEG and LysNTA-Ni^2+^and finally with the E/EC15 fragments by metal affinity binding. d) Field-dependent magnetization curve of the MNPs in organic medium (hexane). e) TGA curves showing the weight loss after organic coating of the MNPs (8 % oleic acid, green curve) and after water transference (18 % MNPs@PMAO (modified with TAMRA), blue curve). f) ζ-potential measurements of MNPs@PMAO (modified with TAMRA) before and after the functionalization with PEG and LysNTA-Ni^2+^. g) Quantification of E/EC15 fragments bound to the MNPs depending on the added amount per mg of Fe, when 120 or 240 μg mg^-1^ Fe were added. For f) and g), black asterisks indicate statistical differences (*p < 0.05; ****p < 0.0001) after a t-Test analysis.

### Synthesis and characterization of MNPs

Zinc-manganese-iron oxide MNPs (Zn_0.29_Mn_0.18_Fe_2.53_O_4_) with a regular octahedral shape were synthesized by a one-pot thermal decomposition method.^49^ The MNPs showed an edge length of 18.6 ± 2.1 nm and a space diagonal of 26.2 ± 1.9 nm, determined from transmission electron microscopy (TEM) images shown in Figure 2b and in Figure S2 (Supporting Information). The synthesized MNPs were stabilized with oleic acid ligands and were soluble in organic solvents (hexane) (Figure 2c). They displayed superparamagnetic behavior and a high saturation magnetization (*M_s_*) value of 99 Am^2^ kg^-1^ (calculated per kg of ferrite) at room temperature (RT) as determined from their corresponding magnetization curve shown in Figure 2d. To use the MNPs in biological media, they were coated with an amphiphilic polymer, poly (maleic anhydride-*alt*-1-octadecene) (PMAO), previously modified with tetramethylrhodamine 5-6-carboxamide (TAMRA) cadaverine, a fluorescent dye to facilitate MNP detection by fluorescence microscopy.^50,51^ After coating with PMAO, the MNPs were analyzed by thermogravimetric analysis (TGA) to quantify the organic coating. TGA revealed a weight contribution of 8% from oleic acid and of 18% from the PMAO coating (Figure 2e).

PMAO provided carboxylic groups, enabling further functionalization with the nitrilotriacetic acid (NTA) derivative Nα,Nα-bis(carboxymethyl)-L-lysine hydrate (LysNTA) and α-methoxy-ω-amino polyethylene glycol (PEG, 5000 Da) as depicted in Figure 2c. This functionalization, necessary for the oriented immobilization of E/EC15 fragments and for improving colloidal stability, respectively, was carried out following our previously reported method.^40^ The resulting nanoparticles MNPs@PMAO@(PEG + LysNTA-Ni^2+^) showed an increase in the hydrodynamic diameter when compared to MNPs@PMAO, as determined by dynamic light scattering (DLS) (Table S1, Supporting Information). Similarly, the MNPs evidenced an increase in the surface charge from -25.3 ± 0.8 mV for MNPs@PMAO to -16.1 ± 0.5 mV for MNPs@PMAO@(PEG + LysNTA-Ni^2+^) (Figure 2f), in accordance with our previous results.^40^ The amount of Ni^2+^ incorporated on the MNP surface was quantified by inductively coupled plasma – atomic emission spectroscopy (ICP-AES), obtaining a ratio of 0.37 μmols of Ni^2+^ per mg of Fe (approximately 2800 LysNTA-Ni^2+^ complexes per MNP). These results confirmed a successful functionalization of the MNPs, allowing further bioconjugation with E/EC15 fragments.

### Bioconjugation of E/EC15 fragments with the MNPs

E/EC15 fragments were then immobilized *via* metal chelation on the MNPs functionalized with LysNTA-Ni^2+^ allowing their orientation *via* their 6His-tag.^53,54^ In our previous study, we demonstrated that the density of E-cadherin fragments on the MNP surface plays a pivotal role in both the functionality of the cadherin fragments and their recognition by cellular cadherins.^40^ To optimize the E/EC15 surface density, MNPs were functionalized with two different fragment amounts (120 or 240 µg mg^-1^ Fe, Figure 2g). At the lower ratio (120 µg mg^-1^ Fe), 72 % of the added protein bound to the MNPs, corresponding to an average of 11 E/EC15 fragments per MNP. Increasing the cadherin amount to 240 µg mg ¹ Fe resulted in a binding efficiency of 53 % and a higher surface coverage, with approximately 17 fragments per MNP.^40^

To assess the functionality of both bioconjugates, an aggregation assay was performed in the presence and absence of calcium ions, following a previously described method.^40,55,56^ As shown in Figure 3a, MNP aggregation occurred after 30 minutes exclusively in the presence of calcium ions, consistent with calcium-dependent homophilic interactions between cadherins present on the surface of distinct MNPs. These aggregates were dissociated when adding the calcium chelator ethylenediaminetetraacetic acid (EDTA) (5 mM), confirming the specificity of the interaction. These results mirror those previously observed for bioconjugates functionalized with fragments containing only two E-cadherin domains (E/EC12).^40^ While both constructs promoted aggregation, MNPs@E/EC15 exhibited faster kinetics, forming aggregates within 30 minutes, whereas MNPs@E/EC12 required at least 12 hours. The bioconjugates were then incubated for 30 minutes with Madin-Darby canine kidney (MDCK) cells, known as a model epithelial cell line that expresses E-cadherin.^57^ As shown in Figure 3b, both bioconjugates were able to recognize cellular cadherins, forming a punctuate pattern at the cells edges that could be observed by fluorescence microscopy. However, MNPs functionalized with 17 E/EC15 fragments per MNP (high density) evidenced a clearer labeling and were therefore selected for further experiments (see Table S1 in Supporting Information for further characterization).

**Figure 3.**
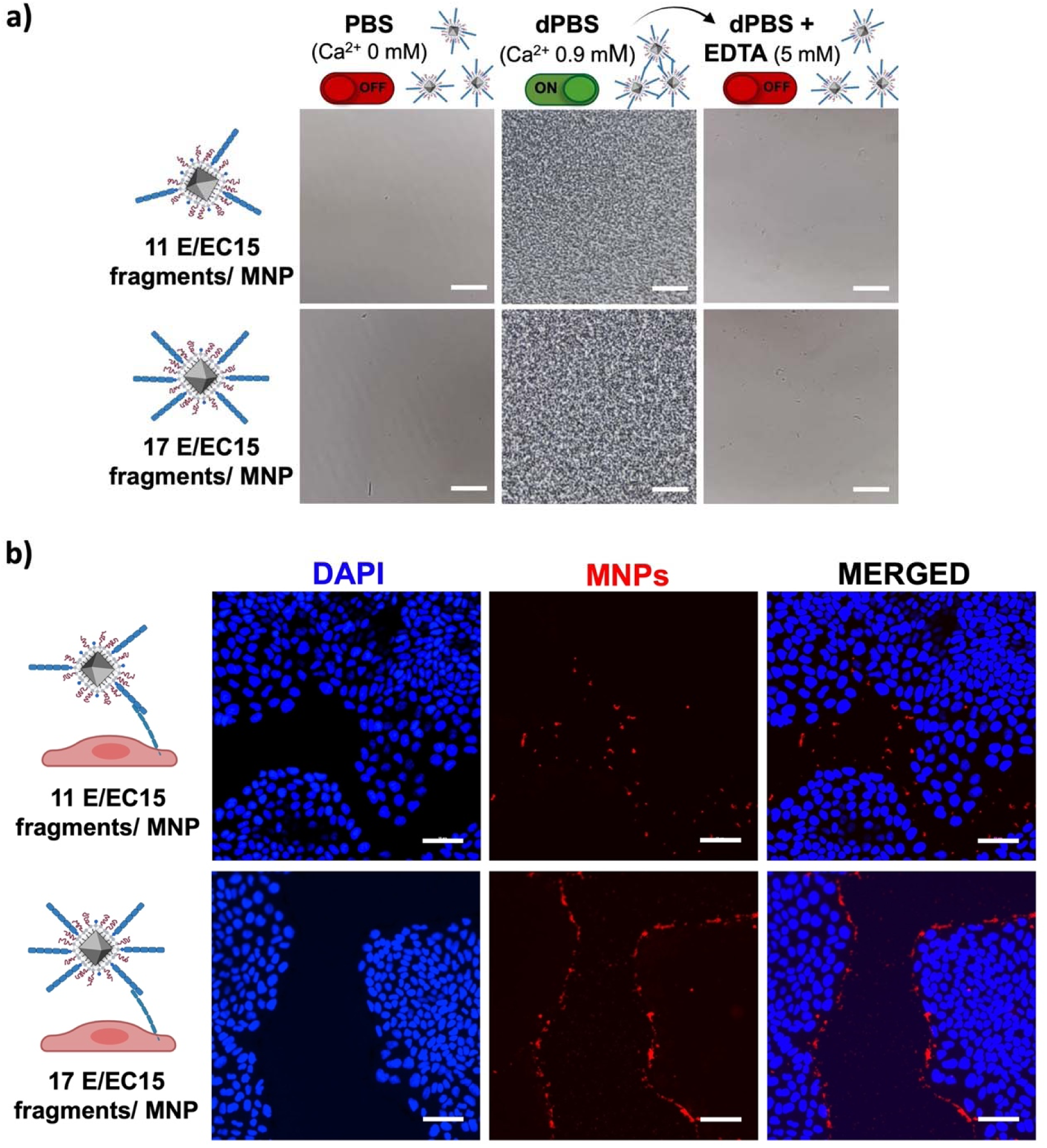
Functional characterization of MNP@E/EC15. a) Aggregation assay in presence and absence of calcium ions using bioconjugates with two different densities of E/EC15 fragments (11 or 17 fragments per MNP). A reversible aggregation was observed upon EDTA (5 mM) addition. Scale bar: 100 μm. b) Fluorescence microscopy images of MDCK cells incubated with the two bioconjugates with different E/EC15 surface density; nuclei were stained with 4′,6-diamidino-2-phenylindole dilactate (DAPI; blue); MNPs containing TAMRA are shown in red. Scale bar: 50 μm.

Thereafter, these bioconjugates were compared with high-density bioconjugates functionalized with E/EC12 fragments (60 fragments per MNP).^40^ MNPs functionalized with E/EC12 fragments were able to label cells; however, bioconjugates containing E/EC15 fragments achieved enhanced labeling efficiency and specificity, with interactions primarily localized on the cell membranes (Figure S3a, Supporting Information).

These results suggest that the reduced accessibility of the two E-cadherin domains (E/EC12) on the MNP surface, determined by the size of the E/EC12 fragment and the thickness of PEG grafting (Figure S3b, Supporting Information), may hinder their trans-interaction with opposing cadherins. Additionally, the presence of a methionine residue as the first amino acid translated in the E/EC12 fragments may also contribute to this reduced interaction.^41,42^ In conclusion, the use of E/EC15 fragments significantly improved the functional MNP aggregation and labeling of MDCK epithelial cells when compared to E/EC12 fragments, confirming the importance of fragment design for cadherin targeting.

Since our goal was to stimulate E-cadherins located on the cell membrane, we assessed the internalization dynamics of the MNPs@E/EC15 in epithelial cells to estimate the post-incubation timeframe in which the bioconjugates remained on the membrane before internalization. To this end, cells were incubated with the bioconjugates for 30 minutes, washed to remove unbound particles, and then fixed 30 minutes or three hours post-treatment. Fixed cells were then imaged by fluorescence microscopy and scanning electron microscopy (SEM) (Figure 4a-b).

**Figure 4.**
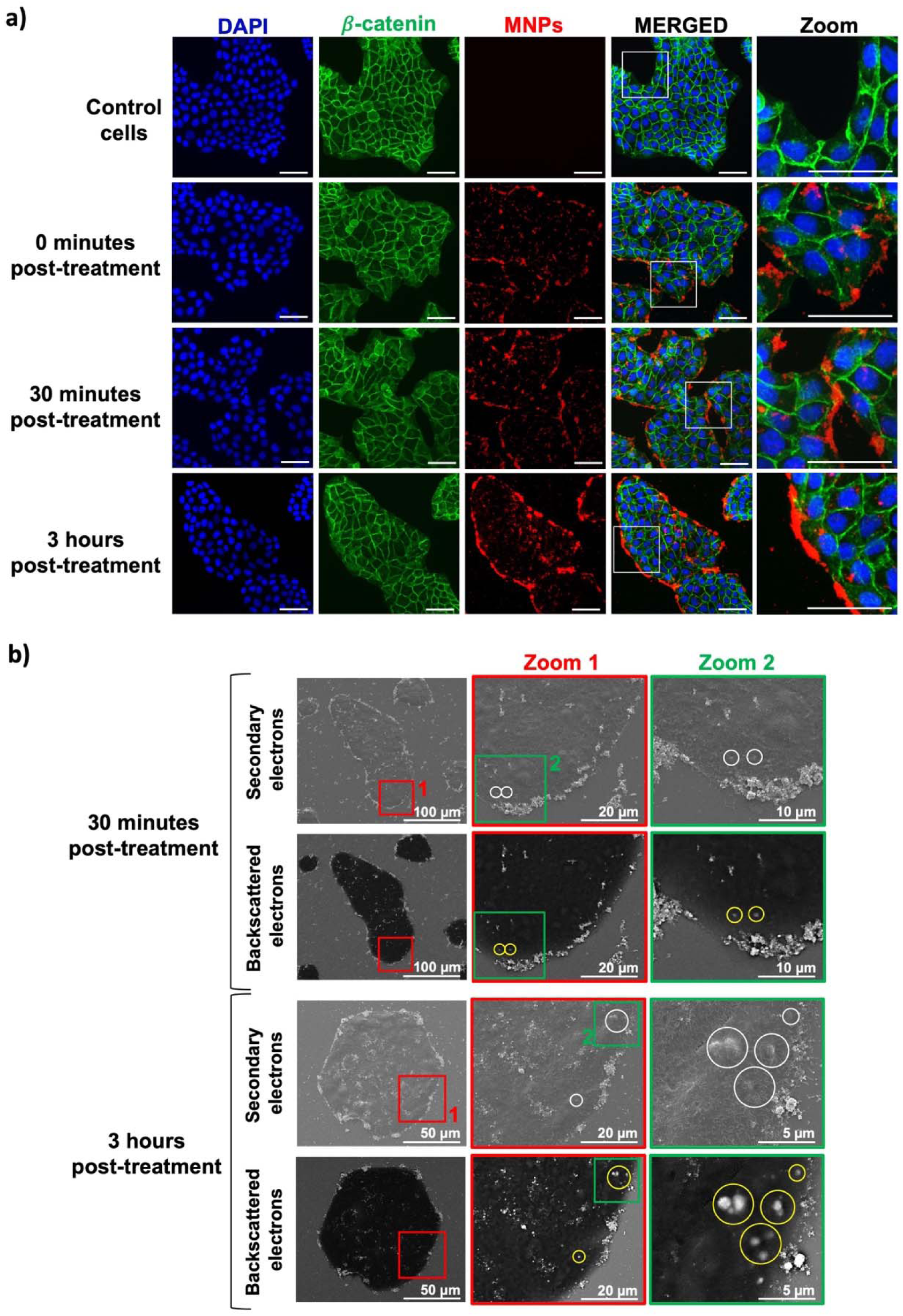
Fluorescence microscopy and SEM imaging of MDCK cells incubated 30 minutes with the bioconjugates, washed and fixed 0, 30 minutes or 3 hours post-treatment. a) Fluorescence microscopy images showing MDCK cells labelled with the bioconjugates; nuclei were stained with DAPI (blue); β-catenin was immunostained and is shown in green (AF488); MNPs containing TAMRA are shown in red; scale bar: 50 μm. b) SEM images of MDCK cells labelled with the bioconjugates. Imaging was performed using secondary and backscattered electrons detectors. In both cases the bioconjugates are shown as bright structures; zones with internalized bioconjugates are pointed out with white circles in secondary electron images and yellow circles in backscattered electrons images where the internalized bioconjugates were easily detected; green and red squares indicate different zoom areas.

For immunofluorescence analysis, the cell membrane was labeled using β-catenin immunostaining, as β-catenin is closely associated with the intracellular domain of E-cadherin and predominantly localizes at the cell edges.^15,25^ As shown in Figure 4a, the bioconjugates (red punctuate pattern) remained mainly located at the edges of the cell colonies even three hours post-treatment, suggesting that no internalization occurred during this time (Figure 4a, Zoom). To discriminate between intra- and extracellular MNP localization, SEM imaging was performed (Figure 4b). Imaging with secondary electrons was performed to track the presence of the MNPs@E/EC15 on the surface of the cell membrane without being internalized (bright structures). On the other side, backscattered electrons capable of visualizing deeper regions of the sample were used to identify internalized bioconjugates (highlighted with yellow circles in Figure 4b). Consistent with the immunofluorescence results, the majority of bioconjugates remained attached on the cell membrane for even three hours post-treatment, indicating minimal internalization. These findings established this timeframe as optimal for subsequent magnetic stimulation experiments.

### Magnetic field applicator

We hypothesized that magnetostimulation of cells with MNPs@E/EC15 interacting with cellular cadherins would exert mechanical stimulation on these biomolecules, potentially promoting β-catenin nuclear translocation and triggering the activation of the canonical Wnt pathway (Figure 1). A customized magnetic stimulation device was used to apply magnetic fields to circular culture dishes, as shown in Figure 5a. The system, partially based on the device previously described by Moreno-Mateos *et al.*,^58^ consisted of four arrays of permanent magnets (10 x 10 x 3 mm thick, NdFeB) positioned at 90° intervals around the dish. These magnet arrays were organized into two opposing pairs, each mounted on an independent motion axis (Figure 5b). The axes operated synchronously and in a cyclic manner: in the first half of the cycle, one pair of opposing magnet arrays moved towards the culture dish while the perpendicular pair simultaneously moved away (position A). In the second half of the cycle, the roles were reversed (position B). This system was configurated to operate at a selected frequency of 0.5 Hz, enabling dynamic and oscillating magnetic stimulation of the sample.^58^

**Figure 5.**
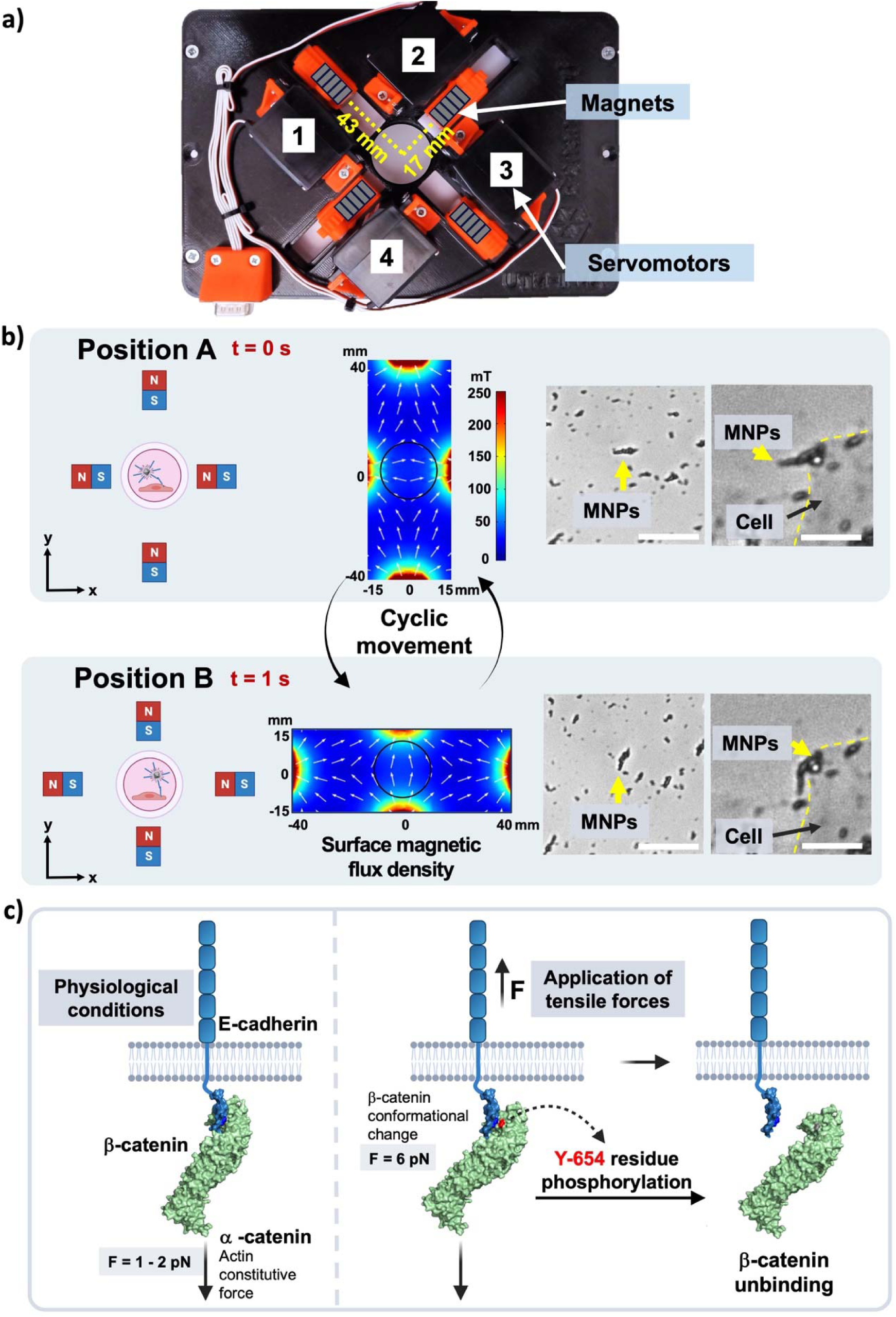
Magnetomechanical stimulation of cellular E-cadherin. a) Image of the magnetic device in which the four arrays of magnets are depicted in gray and the shortest and longest distance from the center of the dish to the magnets are shown in yellow. b) Movement of the magnet arrays consisted of a cyclic pattern alternating between position A – B with a frequency of 0.5 Hz; the magnetic field lines in position A and B were theoretically simulated using COMSOL Multiphysics. Under magnetic field application, the bioconjugates in dPBS or after cell labeling formed chain-like structures, which aligned dynamically with the magnetic field. Cell boundaries are indicated by a yellow dotted line. Scale bar: for bioconjugates in dPBS 25 μm and for cell labeling 10 μm. c) Scheme of the effects promoted by the application of tensile forces on E-cadherin. Under normal physiological conditions, E-cadherin experiments a constitutive force (1-2 pN) exerted by actin.^66^ Upon the application of external tensile forces (6 pN) β-catenin can be phosphorylated and permanently dissociated from E-cadherin.

Magnetic stimulation was performed using five permanent magnets per array. Theoretical simulations of the surface magnetic flux density generated by the whole magnetic applicator were conducted using COMSOL Multiphysics software, evidencing a dynamic rotational magnetic field originated by the magnet movement (Figure 5b). Within the effective culture area of the dish (21 mm in diameter), this configuration generated a low gradient (<15 mT mm^-1^).^49^ Lastly, during the alternating movement of the magnets from position A to B, the magnetic field at the center of the dish ranged between 14 and 33 mT (Figure S4, Supporting Information).

To estimate the theoretical forces exerted by a single MNP, it is important to consider that under a rotational magnetic field MNPs experience a magnetic torque (determined by their intrinsic properties) that tends to align their magnetic moment with the field direction.^59,60^ Theoretical calculations estimated that a single MNP located at the center of the culture dish at maximum field intensity (33 mT) (Figure 5b), exerted a torque of 1.1 x 10^-21^ N m, which corresponds to a force of 80 fN calculated as indicated in our previous work.^49^ However, as shown before, our bioconjugates formed small aggregates in the presence of calcium driven by the interaction between E/EC15 fragments (Figure 3a), and their intrinsic magnetic interaction. These aggregates, composed of multiple MNPs, progressively aligned into chain-like elongated assemblies that exhibited a rotational motion under magnetic stimulation, as shown in Figure 5b and Figure S5 (Supporting Information), which can amplify the mechanical force transmitted to cellular cadherins.^49,61,62^ For instance, Shen *et al.* reported a forces as high as 100 pN for similar MNPs forming assemblies of μm,^63^ while in our previous work an assembly with a length of 11 μm and a width 2.6 μm could generate a theoretically magnetic force of 32 pN.^49^ Moreover, we have previously demonstrated that these MNPs functionalized with an specific antibody could trigger Piezo1 channel opening,^49^ which requires forces in a range of 2-10 pN.^64,65^

As previously mentioned, β-catenin bound to the cytoplasmic domain of E-cadherin undergoes conformational changes in response to mechanical forces of 6 pN, exposing its Y654 phosphorylation site and leading to its irreversible dissociation from E-cadherin.^27^ Given that the forces exerted by the MNP assemblies can fall within this range, we proposed that our system could effectively promote β-catenin dissociation from E-cadherin, thereby triggering the Wnt/β-catenin pathway activation (Figure 5c).

### MNPs@E/EC15 alone or combined with magnetic stimulation affect the transcriptomic landscape of MDCK cells

To investigate the transcriptomic response of MDCK cells to MNPs@E/EC15 bioconjugates, either alone or combined with magnetic stimulation, RNA sequencing was performed. Four experimental conditions were evaluated, as shown in Figure 6a: (i) ‘Control cells’, (ii) ‘Magnet’ (cells magnetostimulated but not incubated with MNPs@E/EC15), (iii) ‘MNPs’ (cells treated with the MNPs@E/EC15), and (iv) ‘MNPs + Magnet’ (cells incubated with MNPs@E/EC15 and magnetostimulated). The experimental workflow involved incubating MDCK cells with the bioconjugates for 30 minutes, followed by magnetic stimulation for three hours and a subsequent 24 hours incubation prior to cell lysis, RNA extraction and RNAseq analysis (Figure 6b). For all conditions, experiments were performed in independent biological triplicates, showing a consistent intragroup behavior (Figure S6a-b, Supporting Information).^67^ Differentially expressed genes (DEGs) in each condition compared to the control were identified using a |Log2FC| ≥ 0.5 threshold (Figure S6c, Supporting Information). This value was selected to include both robust and moderate changes in gene expression, thereby preserving subtle yet potentially biologically meaningful regulatory effects. The overlap and distribution of DEGs across the three ‘treatment vs. control’ comparisons are illustrated in the Venn diagram in Figure 6c, which highlights both shared and unique gene expression changes. The cellular response to MNPs@E/EC15, either alone or in combination with magnetic stimulation, was more pronounced than the one elicited by magnetostimulation alone.

**Figure 6.**
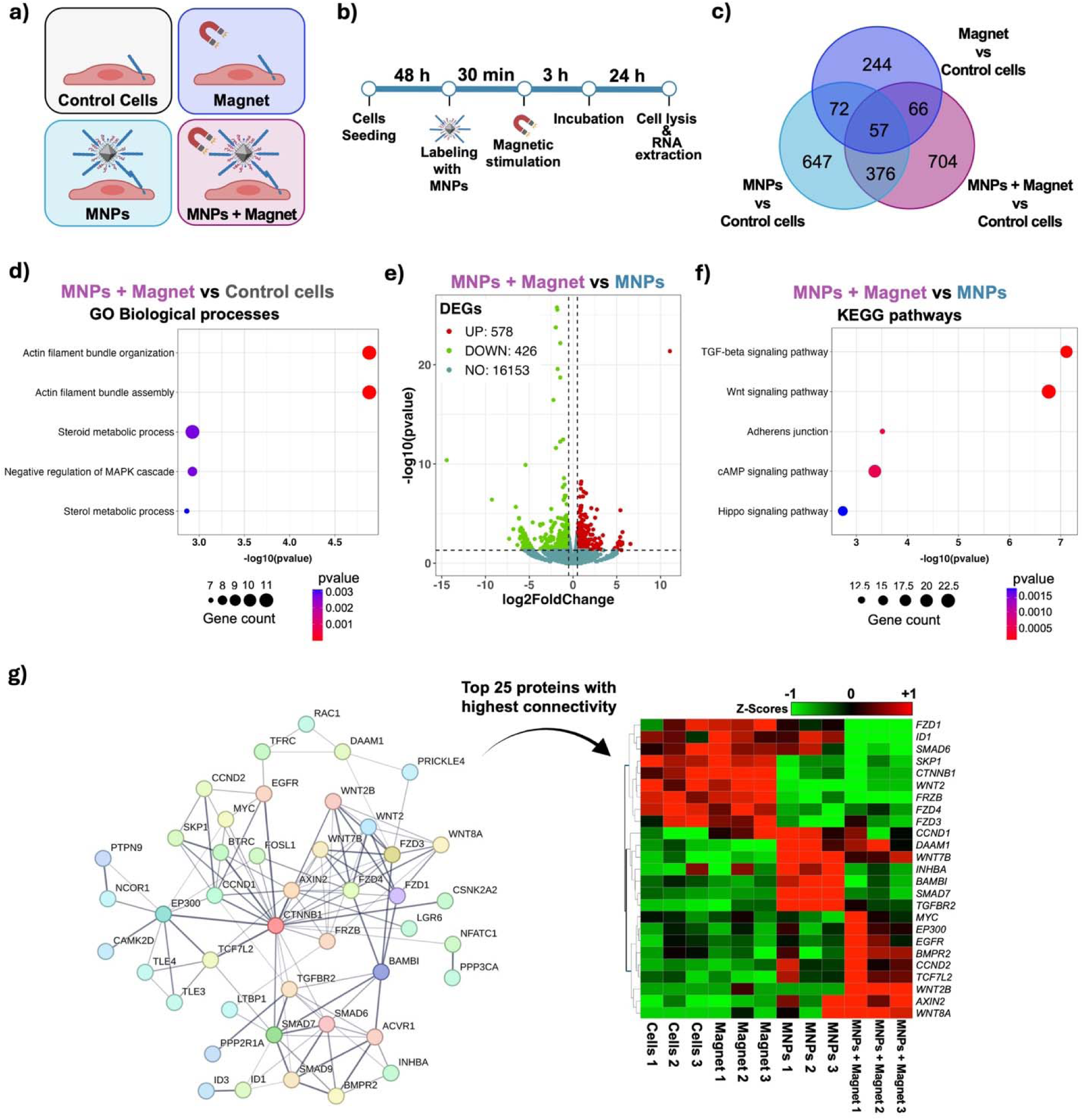
Transcriptomic response of MDCK cells to E-cadherin bioconjugates. a) Scheme of the experimental conditions evaluated in the transcriptomic analysis. b) Timeline of the experiment performed before cell lysis for RNA sequencing. c) Venn diagram showing the number of DEGs exclusively associated to the different treatments versus control comparisons. d) GO enrichment analysis (biological process category) of the 704 DEGs identified in the Venn diagram as exclusively activated in presence of the bioconjugates in combination with magnetic stimulation showing the top 5 terms enriched. e) Vulcano plot showing the 1004 DEGs identified from ‘MNPs + Magnet’ vs MNPs comparison. f) KEGG pathway enrichment analysis of the 1004 DEGs showing the top 5 pathways enriched. g) Protein-protein interaction (PPI) network showing DEGs of Wnt pathway from ‘MNPs + Magnet’ vs ‘Control cells’, and ‘MNPs + Magnet’ vs MNPs comparisons; a heatmap colorized following Z-scores was generated showing the top 25 proteins (Wnt pathway) with more nodes from PPI network.

Among the 647 DEGs uniquely associated with treatment using MNPs@E/EC15 bioconjugates, functional enrichment analysis revealed that the bioconjugates induced a broad regulatory response in cells (Figure S7a-b, Supporting Information). Gene Ontology (GO) analysis indicated a significant enrichment in the positive regulation of developmental processes, a biological process (BP) involving DEGs linked to cell differentiation, organ formation and morphogenesis.^68^ Notably, Wnt signaling emerged as the most enriched pathway (Figure S7c, Supporting Information), suggesting that engagement of E-cadherin by the bioconjugates may influence Wnt signaling.

Although E-cadherin engagement can occur independently of mechanical force, that is, promoted only by the interaction of bioconjugates with cells, other processes such as actin assembly and organization are typically triggered in response to force application on E-cadherin.^69,70^ Consistently, functional analysis of the 704 DEGs identified under combined bioconjugate treatment and magnetic stimulation (‘MNPs + Magnet’ vs ‘Control cells’) (Figure 6d) revealed significant enrichment in actin filament bundle organization and assembly (Figure S8a-b, Supporting Information). Our analysis showed upregulation of key genes involved in cytoskeleton regulation mediated by E-cadherin mechanotransduction, including *RAC1, RHOA, AMOTL2, WAVE2,* and *MYO1C*.^70–73^ Previous studies have demonstrated that E-cadherin mechanotransduction (E-cadherin stiffening, vinculin recruitment, and reinforcement of E-cadherin-actin linkages) can be triggered by mechanical forces applied through magnetic microbeads functionalized with E-cadherin fragments or antibodies.^34^ However, the large size of these microparticles can lead to unintended cytoskeletal responses mainly due to membrane deformation. In contrast, Seo *et al.* demonstrated that using MNPs instead of microbeads minimizes such membrane perturbations, preventing off-target effects and ensuring that cytoskeletal responses are exclusively triggered by force application.^11^ Our findings are consistent with this observation, and provide clear evidence that magnetomechanical stimulation exerted on our MNPs@E/EC15 bioconjugates (‘MNPs + Magnet’) was strong enough to trigger an E-cadherin-mediated mechanotransduction response promoting cytoskeleton reorganization.

While this initial transcriptomic analysis provided information regarding the effects exerted by the bioconjugates alone and in combination with magnetic stimulation, a direct comparison between both treatments (‘MNPs + Magnet’ vs MNPs) was performed to fully discriminate genes exclusively regulated by magnetomechanical stimulation. This comparison revealed 1004 DEGs (Figure 6e and Figure S9a, Supporting Information), which were further subjected to functional enrichment analysis. GO analysis revealed enrichment of molecular functions related to Wnt/β-catenin, mitogen-activated protein kinase (MAPK) or transforming growth factor-beta (TGF-β) pathways. These pathways may contribute to more complex biological processes such as organ morphogenesis or epithelium development (Figure S9b-c, Supporting Information),^74–76^ in which E-cadherin is also directly involved.^73,77^ Consistently, TGF-β and Wnt signaling appeared as the most enriched pathways (Figure 6f), evidencing that Wnt signaling pathway modulation not only occurred by E-cadherin engagement, but also under the combined action of the bioconjugates with magnetic stimulation. Both pathways are known to crosstalk at multiple levels through shared protein mediators including SMAD7, BAMBI or TCF7L2 (all of them also affected by our bioconjugates),^16,78,79^ and are involved in the regulation of key cellular processes such as proliferation, development, wound healing and epithelial to mesenchymal transition (EMT).^15,76^ Several canonical Wnt target genes such as *CTNNB1* (β-catenin), *AXIN2*, *EGFR*, *MYC* and *TCF7L2* were upregulated, all of which are well-known downstream effectors of β-catenin nuclear translocation.^80,81^ To the best of our knowledge, this is the first report demonstrating the remote activation of TGF-β and Wnt signaling pathways *via* E-cadherin-mediated magnetomechanical stimulation using MNPs.

Notably, the Hippo pathway, another well-known mechanosensitive signaling cascade associated with cell proliferation, also ranked among the most enriched pathways in this analysis (Figure 6f). Concordantly, upon biaxial cell stretching the mechanical strain transduced by E-cadherin stimulation has shown to promote YAP and β-catenin activation to drive cell cycle re-entry.^28^ These results support the notion that magnetomechanical stimulation *via* MNPs@E/EC15 bioconjugates effectively modulate multiple force-responsive signaling networks in epithelial cells.

To gain a more in-depth understanding of the Wnt pathway activation induced by magnetomechanical stimulation, we isolated the DEGs associated with Wnt signaling pathway from two comparisons: (i) ‘MNPs + Magnet’ vs ‘Control cells’ and (ii) ‘MNPs + Magnet’ vs MNPs. These genes were visualized using STRING (https://string-db.org) to conduct a protein-protein interaction (PPI) network, which is depicted in Figure 6g. The 25 proteins with the highest connectivity within the network were further visualized in a heatmap to highlight expression trends. Our analysis revealed distinct expression patterns of central players in the Wnt/β-catenin signaling pathway: for instance, *CTNNB1* (β-catenin) was downregulated, while *AXIN2* was upregulated by the bioconjugates, regardless of magnetic stimulation. This reciprocal regulation reflects a well-known negative feedback loop, in which *AXIN2* is activated by β-catenin, but once expressed, *AXIN2* promotes β-catenin degradation.^80^ Additionally, genes such as *SMAD7* and *BAMBI*, considered inhibitors and crosstalk mediators between the Wnt and TGF-β pathways, were upregulated by the bioconjugates but returned to basal expression levels upon magnetic stimulation, at least in the time window in which the transcriptomic analysis was carried out. Notably, *TCF7L2, MYC,* and *EGFR,* key Wnt/β-catenin target genes closely associated with cell proliferation, were upregulated only in response to the combined action of bioconjugates and magnetic stimulation, suggesting a synergistic effect. To validate these transcriptomic findings, we selected *MYC*, *EGFR*, and *SMAD7* for real-time quantitative polymerase chain reaction (RT-qPCR) analysis. The resulting relative mRNA expression levels (2^-ΔΔCt) were consistent with RNA-seq results (Figure S10, Supporting Information), thus supporting the robustness of the transcriptomic analysis.

### Magnetomechanical stimulation with MNPs@E/EC15 bioconjugates promotes Wnt pathway activation and drives cell proliferation

To confirm whether the observed transcriptomic activation of the Wnt pathway translated into functional protein-level changes, MDCK cells were genetically engineered using a lentiviral vector encoding a luciferase reporter under the control of a Wnt-responsive promoter (7xTCF) as depicted in Figure 7a.^82^ Similar surface expression of E-cadherin relative to the parental line was verified by flow cytometry (Figure S11, Supporting Information). In this system, nuclear translocation of β-catenin activates the TCF/LEF promoter, leading to luciferase expression. Upon addition of the luciferase substrate, the resulting luminescence signal constitutes a quantitative measure of the β-catenin-dependent transcriptional activity, which can be quantified spectrophotometrically. Luciferase activity was measured across all four experimental conditions and normalized to total protein content (TPC) to account for variations in cell number (Figure 7b).

**Figure 7.**
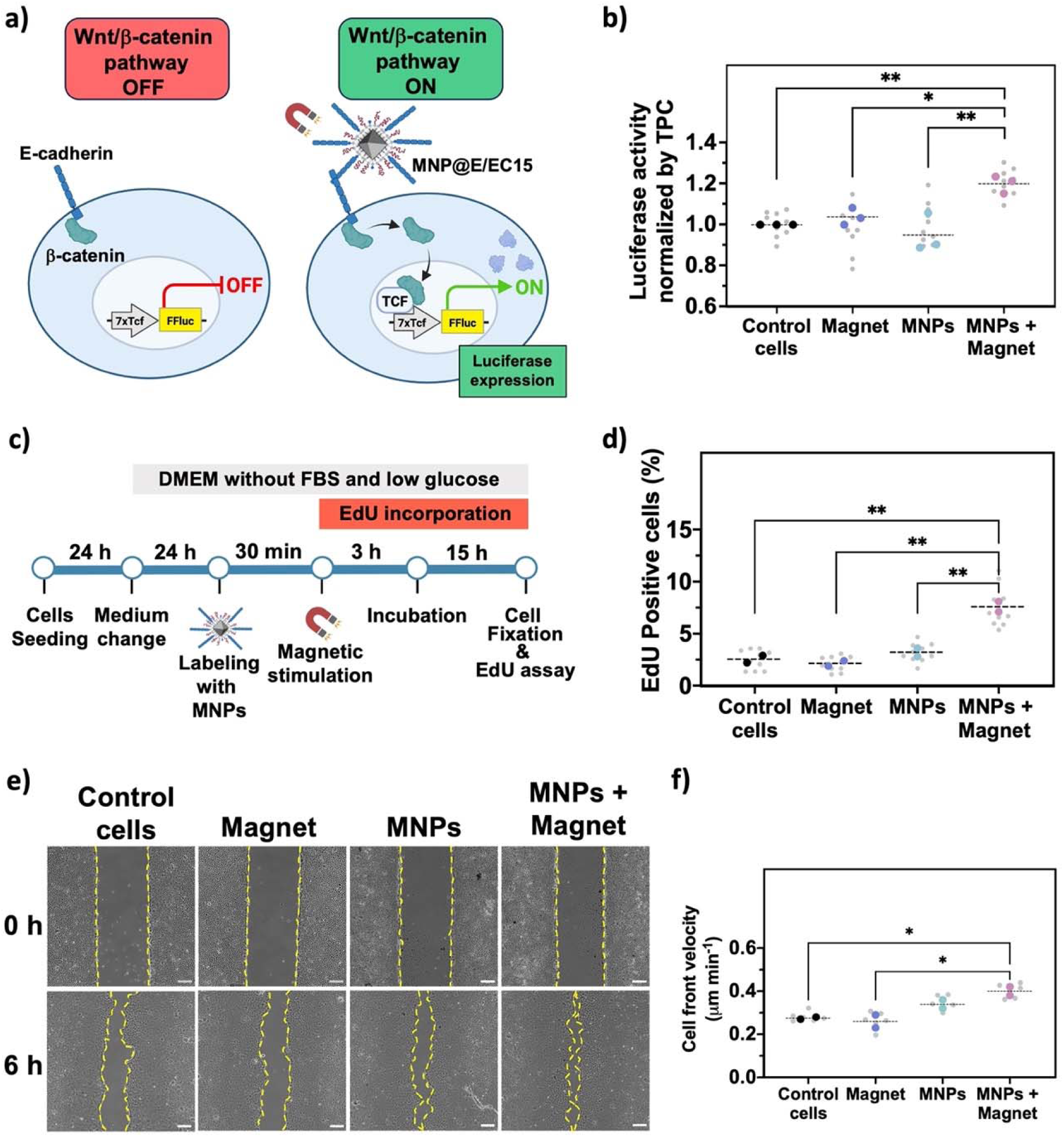
Magnetomechanical stimulation triggers Wnt/β-catenin activation and cellular proliferation. a) Scheme of MDCK cells modified by lentiviral transduction to express a luciferase reporter gene (FFluc) under the control of a promoter activated by β-catenin-TCF complex. Image created in https://BioRender.com. b) Luciferase activity normalized to TPC, showing an increase in the samples treated with bioconjugates combined with magnetic stimulation. c) Timeline of the EdU staining assay. Cells were incubated with EdU reagent for 15 hours before fixing. d) Percentage of EdU-positive cells 15 hours post-stimulation. e) Representative images of the wound healing assay showing cells at 0- and 6-hours post-stimulation. Scale bar: 100 μm. f) Kinetics of gap closure. The slope from gap closure kinetics (Figure S13, supporting information) from 0 – 6 hours post-stimulation was used to calculate the corresponding cell front velocity (μm min^-1^) for statistical analysis following the equation presented in the Supporting Information. Black asterisks indicate statistical differences (*p < 0.05; **p < 0.01; ***p < 0.001; ****p < 0.0001). One-way ANOVA followed by Tukey’s multiple comparison test were performed. Each gray dot corresponds to one technical replicate (3 – 4), and the colored dots corresponds to the mean of each independent experiment performed in different days

As shown in Figure 7b, neither magnetic stimulation alone nor incubation with MNPs@E/EC15 bioconjugates significantly increased luciferase activity compared to ‘Control cells’. However, the combination of bioconjugates with magnetic stimulation resulted in a statistically significant 20% increase in β-catenin-dependent transcriptional activity. This result suggested that although E-cadherin engagement can influence Wnt pathway (Figure S8c, Supporting Information), magnetomechanical stimulation was required to promote Wnt/β-catenin signaling pathway activation. This observation is consistent with findings by Benham-Pyle *et al.* who reported a 20 – 30 % increase in β-catenin transcriptional activity between 6- and 16-hours following mechanical stretching of MDCK cells, followed by a gradual decrease to basal levels.^28^ However, in this study, cell stretching was applied to a dense monolayer of cells without any spatial control, potentially activating additional mechanosensitive proteins beyond E-cadherin. In contrast, our approach enables precise targeting of E-cadherin using nanoscale bioconjugates, demonstrating the ability to exert mechanical rotational forces (torque) on cellular E-cadherins with high spatiotemporal resolution. Notably, Wnt/β-catenin signaling pathway activation was achieved despite MDCK cells forming tightly packed colonies, highlighting the potential of the MNP bioconjugates for the selective stimulation of membrane proteins in complex epithelial architectures.

To determine whether the observed activation of the Wnt/β-catenin signaling pathway translated into functional biological outcomes, we assessed cell proliferation and migration. Proliferation was evaluated using Click-iT™ EdU (5-ethynyl-2’-deoxyuridine) staining, while collective cell proliferation/migration dynamics were analyzed *via* wound healing assays.

EdU is a thymidine nucleoside analog incorporated into newly synthesized DNA during active proliferation (S-phase of the cell cycle). Following incorporation into cells, a fluorophore-labeled azide reacts with the incorporated ethynyl-labeled nucleoside *via* click chemistry, allowing detection of proliferating cells by fluorescence microscopy (Figure S12, Supporting Information). As depicted in Figure 7c, cells were first cultured in low-glucose, serum-free Dulbecco’s Modified Eagle Medium (DMEM) to induce quiescence and synchronize proliferation rates prior to the incubation with the bioconjugates. In Figure 7d, the number of EdU-positive cells (proliferating cells) for each condition is shown. ‘Control cells’, as well as ‘Magnet’ and ‘MNPs’ treatments, exhibited low proliferation rates (2.0–3.2 % EdU-positive cells). By contrast, bioconjugates combined with magnetic stimulation (‘MNPs+Magnet’) significantly increase proliferation to 7.6 % EdU-positive cells, representing a threefold rise over the control. This proliferative response agrees with the findings of Benham-Pyle *et al.,* who evidenced a similar increase in EdU-positive cells 24 hours after mechanical stretching, further supporting the role of mechanical stimulation in enhancing proliferation.^28^

A similar trend was observed in the wound healing assay. As shown in Figure 7e-f, the most pronounced effect was observed when magnetic stimulation was applied in combination with the bioconjugates. Only under this condition wound closure rate became significantly faster than in ‘Control cells’. While the increased proliferation observed in the EdU assay likely contributed to the faster wound closure, cell migration dynamics could also play a role. Mechanical forces experienced by cells through E-cadherin during collective migration are known to influence this process,^83^ but further investigation is required to support the contribution of migration in the context of our experiments.

Overall, the results from both the EdU incorporation and wound healing assay strongly suggest that magnetomechanical stimulation of cellular E-cadherin *via* MNPs@E/EC15 bioconjugates enhance epithelial cell proliferation rate. Previous studies have attempted to activate the canonical Wnt pathway by targeting and stimulating its main associated receptor (Frizzled) using MNP clusters (250 nm) functionalized with either anti-Frizzled antibodies or Wnt-derived peptides.^84–86^ However, in these approaches, it remains unclear if Frizzled receptor activation occurs through a mechanotransduction process, instead suggesting a clustering mechanism that leads to β-catenin nuclear translocation. Moreover, the use of high-strength magnetic fields (120 mT) and compressive forces arising from the configuration of the magnetic applicator (magnets located under the sample), may trigger non-specific membrane responses upon particle recognition, such as premature endocytosis. In contrast, our approach uses E-cadherin bioconjugates to apply mechanical rotational forces (torque) directly through E-cadherin. This previously unexplored strategy demonstrates that E-cadherin magnetomechanical stimulation can modulate proliferative signaling by activating the Wnt pathway, but also by influencing other pathways such as TGF-β or Hippo rather than acting through a single cascade (Figure 8).

**Figure 8:**
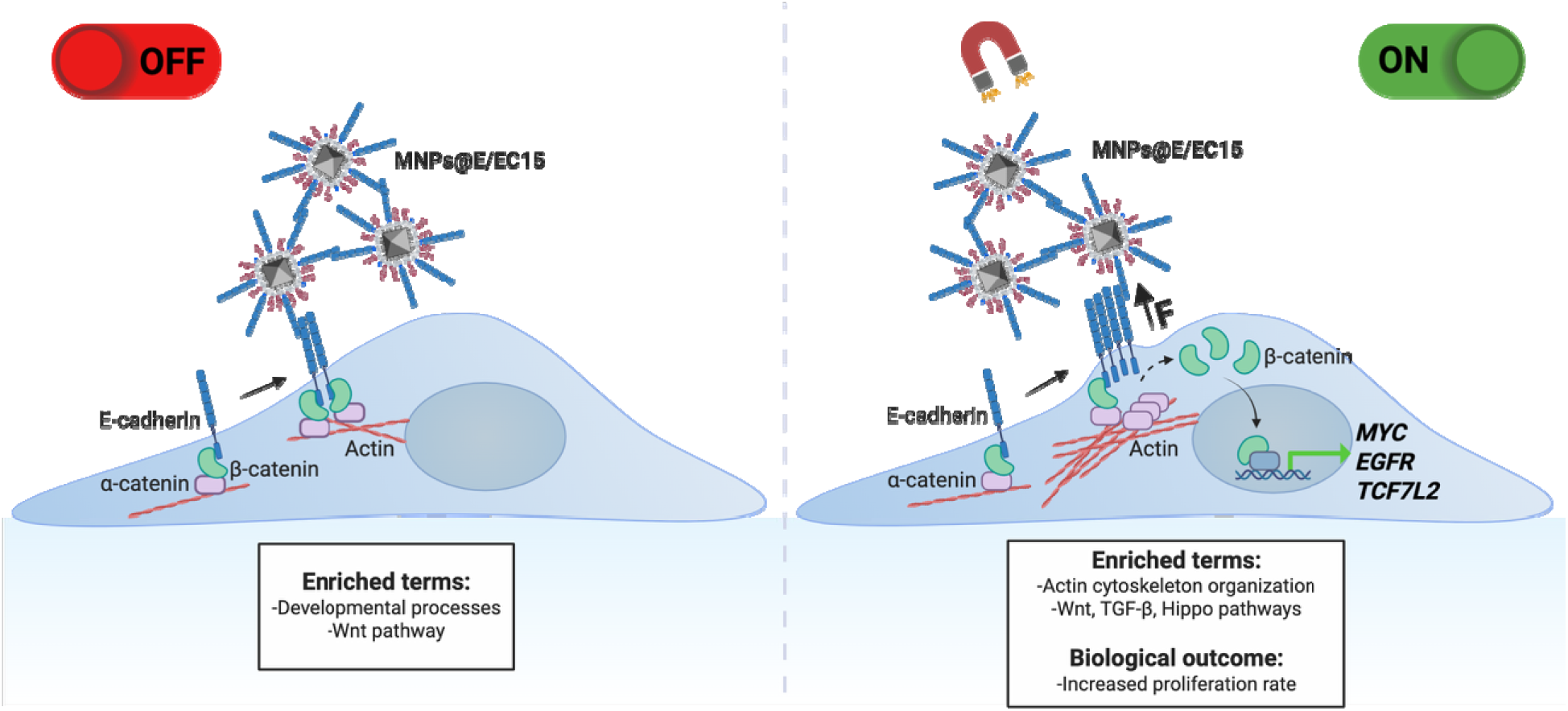
Overview of the effects observed in MDCK cells after the treatment with the MNPs@E/EC15 bioconjugates alone or when combined with magnetic stimulation. Cellular E-cadherin interaction with the bioconjugates potentially leads to E-cadherin engagement which in turn modulates Wnt pathway. Conversely, the application of magnetomechanical forces on E-cadherins promoted the activation of genes related with actin organization as well as an increment of β-catenin translocation leading to the expression of Wnt target genes but also affecting TGF-β and Hippo pathways. Image created in https://BioRender.com.

## Conclusions

In this study we developed a novel magnetogenetic tool to modulate the Wnt/β-catenin pathway with high precision, based on a mechanotransduction process.

To do so, we synthesized E-cadherin-functionalized MNPs (MNPs@E/EC15) and demonstrated their ability to engage with cellular cadherins and transduce mechanical signals with high specificity and spatial control. By tuning the surface density of E-cadherin fragments, we optimized the bioconjugates for effective cell membrane binding with minimal internalization. This efficient cell labeling allowed us to mechanically stimulate the cellular E-cadherins by the application of a weak magnetic field (33 mT), being able to induce enough force to promote E-cadherin mediated responses.

Transcriptomic analysis revealed that MNPs@E/EC15 alone induced significant gene expression changes, including the enrichment of Wnt pathway. When combined with magnetomechanical stimulation, the bioconjugates triggered a distinct mechanotransduction response, activating Wnt/β-catenin and additional signaling pathways such as actin cytoskeleton organization, and TGF-β signaling. Wnt/β-catenin activation was supported by protein-level validation using a Wnt-responsive luciferase reporter system, which confirmed increased β-catenin transcriptional activity upon magnetomechanical stimulation of the bioconjugates (‘MNPs + Magnet’). This supports the hypothesis that mechanical forces applied to cellular E-cadherins through the bioconjugates are sufficient to induce β-catenin nuclear translocation and Wnt pathway activation.

Functionally, these molecular changes translated into increased proliferation and accelerated wound closure, as demonstrated by EdU proliferation assays and wound healing experiments. The significant rise in EdU-positive cells and the faster gap closure in bioconjugate-treated cells under magnetic stimulation suggest a direct influence on cell cycle progression.

Overall, this work establishes MNPs@E/EC15 as a powerful magnetogenetic tool for probing E-cadherin mechanobiology and demonstrates their potential for precise, non-invasive modulation of cell proliferation in epithelial systems, surpassing other methods that lack spatial control. Unlike magnetic tweezers, this tool can activate many cells at the same time and could be potentially translated *in vivo* due to the low magnetic fields that are required. These findings establish a new paradigm for remotely modulating Wnt pathway through E-cadherins, opening new avenues for studying mechanotransduction and developing targeted therapeutic strategies for regenerative medicine and tissue engineering based on mechanical stimulation.

### Experimental Section Reagents and materials

All commercially available reagents were used as supplied unless otherwise stated. Sodium chloride, sodium tetraborate decahydrate, sodium cacodylate, glutaraldehyde, boric acid, Na_2_HPO_4_, NaH_2_PO_4_, iron(III) acetylacetonate, manganese(II) acetylacetonate, zinc(II) acetylacetonate, oleic acid (OA), 1,2-hexadecanediol, PMAO (MW: 30000–50000 g mol^-1^), 1,2-dihydroxybenzene-3,5-disulfonic acid (Tiron), N_α_,N_α_-bis(carboxymethyl)-L-lysine hydrate (LysNTA), nickel(II) chloride hexahydrate, Fluorsave, and Gel-Red were purchased from Sigma-Aldrich. α-methoxy-ω-amino poly(ethylene glycol) (PEG, MW: 5000 Da) were purchased from Rapp Polymere GmbH. Imidazole, benzyl ether, hexane, chloroform stabilized with ethanol, 1-ethyl-3-(3-dimethyl aminopropyl)carbodiimide (EDC), Bradford assay reagent, bovine serum albumin standard (BSA), Pierce™ BCA Protein Assay Kit (Reference: 23225), Lipofectamine 3000 transfection reagent (Reference: L3000002), ultrapure agarose, Click-iT EdU cell proliferation kit for imaging (Reference: C10338), Alexa Fluor 555 dye, High-Capacity cDNA Reverse Transcription Kit (Reference: 4368814) were obtained from Thermo Scientific. Absolute ethanol, sodium hydroxide (NaOH), hydrochloric acid (HCl), nitric acid (HNO_3_), ethylenediaminetetraacetic acid (EDTA), Triton X-100, and tris(hydroxymethyl)aminomethane (Tris) were obtained from Panreac. HisTrap HP His tag protein purification columns (Reference: 17524801) were purchased from Cytiva. Tetramethylrhodamine 5- and 6-carboxamide (TAMRA) cadaverine was obtained from AnaSpec. Amicon centrifugal filter units (100 kDa MWCO) and 0.22 μm pore size 13 mm diameter cellulose acetate membrane filters were obtained from Millipore. Four compartments µ-Dish 35 mm (Reference:80416) and culture insert 2-well were obtained from ibidi. Total RNA purification kit (Reference: 17200) and DNAse-I were obtained from Norgen Biotek Corp. NZYSupreme qPCR Green Master Mix (Reference: MB41903) was obtained from NZYTech. 4–15 % Mini-PROTEAN TGX precast protein gels were purchased from Bio-Rad. DMEM, Optimal Modified Eagle Medium (Opti-MEM), fetal bovine serum (FBS), 1X glutaMAX, phosphate buffered saline (without Ca^2+^ and Mg^2+^) (PBS), Dulbecco’s phosphate buffered saline (with Ca^2+^ and Mg^2+^) (dPBS), and antibiotics for cell culture (penicillin-streptomycin, 10000 U mL^-1^) were obtained from Gibco. Paraformaldehyde was purchased from Electron Microscopy Sciences. Purified mouse anti-β-catenin antibody (Reference: 610153) was obtained from BD Biosciences. Goat Anti-mouse Alexa Fluor 488 (Reference: A-11001), and 4′,6-diamidino-2-phenylindole dilactate (DAPI) were obtained from Invitrogen. Luciferase assay system kit (Reference: 1500) and Trypsin Gold mass spectrometry grade were obtained from Promega. Buffers were prepared according to standard laboratory procedures. Milli-Q water (resistivity of 18.2 MΩ cm^-1^ at 25 °C) was obtained using a Milli-Q Advantage A10 system.

### E/EC15 Expression and Purification

The recombinant E-cadherin fragment, comprising all five extracellular cadherin domains (E/EC15) fused to a C-terminal 6His tag, was expressed in HEK-293 cells stably transduced. The plasmid used encoded nucleotides corresponding to amino acids 1-669 of mouse E-cadherin, including the N-terminal prodomain with its signal peptide, and the mentioned His tag. The cells were kindly provided by Dr. Helene Feracci (CNRS, Bordeaux, France).^44^ Thus, the cells produced the full-length precursor protein, processed the prodomain naturally, and secreted the mature E/EC15 fragment into the extracellular medium, facilitating its subsequent purification. Cells were grown to 80-90 % confluence in DMEM, after which the medium was replaced with Opti-MEM, and cells were incubated for five days to allow E-cadherin expression and secretion. Afterwards, E-cadherin-containing medium was collected and centrifuged for 10 min at 2000 xg to remove cellular debris, and the supernatant was stored at 4 °C until purification.

The expressed E-cadherin was purified from the collected supernatants by metal affinity chromatography using a Hi-Trap column charged with Ni^2+^ ions for binding His-tagged proteins, on an AKTA chromatography system (Amersham Biosciences). The collected medium was mixed with binding buffer (BB) consisting of 20 mM sodium phosphate (Na_2_HPO_4_ + NaH_2_PO_4_) and 0.5 M NaCl in a 2:1 ratio, fixing a final concentration of 30 mM imidazole after mixing to avoid non-specific binding. The column was equilibrated with BB, after which the mixture containing the medium was loaded. Non-specific proteins were washed with washing buffer (similar to BB containing 30 mM imidazole), and the E/EC15 fragments were eluted with elution buffer consisting of 20 mM sodium phosphate (Na_2_HPO_4_ + NaH_2_PO_4_), 0.5 M NaCl and 500 mM imidazole.

### Protein Identification by Peptide Mass Fingerprinting and MS/MS Analysis

Protein identification was performed by analyzing the peptide mass fingerprint (MS) and the fragmentation spectra (MS/MS) of peptides generated after trypsin digestion. First, the fraction collected from AKTA purifier (5 µL) were run in an SDS-PAGE (4-15 % gradient) at 120 volts for one hour. Proteomic analyses were performed in the Proteomics Platform of “Servicios Científico Técnicos del CIBA (IACS-Universidad de Zaragoza)”. Briefly, digestion of the polyacrylamide gel band was carried out in an automatic digester (Intavis, Bioanalytical Instruments). The gel band was sequentially washed with water, 100 mM ammonium bicarbonate, and acetonitrile (ACN). Next, the sample was reduced by incubation with dithiothreitol (10 mM) at 60 °C for 45 min and alkylated by incubation with iodoacetamide (50 mM) at RT for 30 min in the dark. Finally, proteins were digested with trypsin overnight at 37°C (5 ng μL^-1^). Digestion was stopped by adding 0.5 % trifluoroacetic acid, and tryptic peptides were extracted sequentially using increasing concentrations of ACN in water. The samples were evaporated in a concentrator and resuspended in 2 % ACN with 0.1 % formic acid.

Protein identification was performed using a hybrid triple quadrupole/linear ion trap mass spectrometer (6500QTRAP+, Sciex) coupled to a nano/micro-HPLC system (Eksigent LC425, Sciex). The search engine used for protein identification was MASCOT (MatrixScience) with public protein sequence databases (SwissProt, NCBI).

### Synthesis of Zinc-Manganese-Iron Oxide Nanoparticles

In brief, 3.33 mmol of Fe(acac)_3_, 1 mmol of Mn (acac)_2_, and 0.67 mmol of Zn(acac)_2_ were placed in a three-neck flask with 5.29 mL of oleic acid as surfactant and 2.58 g of 1,2-hexadecanediol as reducing agent in 50 mL of benzyl ether. The mixture was mixed and mechanically stirred at 100 rpm under vacuum for 30 min at RT and afterwards heated to 200 °C for two hours (at a rate of 5 °C min^-1^), and then heated up to 285 °C (at a rate of 3 °C min^-1^) for another 2 hours, all under a flow of nitrogen. Finally, the mixture was cooled to RT by removing the heat source. Under ambient conditions, an ethanol excess was added to the mixture, and a black material was precipitated and separated with a permanent magnet. The product was suspended in hexane, then precipitated with ethanol. This suspension-precipitation cycle was repeated four times, resulting in a brownish-black hexane dispersion stored at 4 °C until further use.

### Coating of MNPs with poly(maleic anhydride-*alt*-1-octadecene) (PMAO)^50^

225 mg of PMAO (MW: 30000-50000 g mol^-1^) were dissolved in 18 mL of CHCl_3_. At the same time, 2 mg of TAMRA-cadaverine were dissolved in absolute ethanol (1 mg mL^-1^) and transferred to the flask containing the PMAO solution, leaving the mixture protected from light and under magnetic stirring at RT overnight. 180 mL of CHCl_3_ were added to the mixture, and then 5 mg of Fe of the hydrophobic nanoparticles (previously washed once with ethanol and resuspended in 2 mL of CHCl_3_) were added dropwise in an ultrasonic bath. The mixture was kept in the bath for 30 min at RT. The excess of CHCl_3_ was evaporated using a rotary evaporator at a pressure of 200 mbar at 40 °C until a final volume of 5-10 mL. Once the organic solvent was eliminated, 20 mL of 0.05 N NaOH were added to the mixture to hydrolyze the anhydride groups present in the PMAO and to confer stability in water to the MNPs. The remaining organic solvent was evaporated as before, this time under a pressure of 200 mbar and at 70 °C until a final volume of 10-20 mL. The excess of PMAO was eliminated by four ultracentrifugation steps at 70000 xg per 2 h each one, and the resultant nanoparticle suspension MNPs@PMAO was filtered using a Millipore filter (0.22 μm) to remove MNP aggregates and stored at 4 °C protected from light.

### Determination of Iron Concentration

After each coating, functionalization, or bioconjugation step, determination of iron concentration was performed.^87^ First, 5 μL of MNPs were diluted in 45 μL of solvent (hexane or water) and digested with 100 μL of aqua regia solution (HCl:HNO_3_; 3:1) at 60 °C for 15 min. Then, the samples were diluted up to 500 μL, and 50 μL were used for the iron quantification by mixing the digested samples with 0.25 M 1,2-dihydroxybenzene-3,5-disulfonic acid (Tiron), a molecule that forms a colored complex with iron and can be measured by spectrophotometry (480 nm).^88^ The samples were measured on a Biotek Synergy H1 UV/vis microplate spectrophotometer and compared with a standard calibration curve obtained with solutions of known iron concentrations (0, 100, 200, 400, 600, 800 μg Fe mL^-1^). All iron concentration determinations were carried out in triplicate.

### Functionalization of MNPs@PMAO-TAMRA with PEG, LysNTA-Ni^2+^ and E/EC15^40^

As linker for the His-tagged E/EC15 fragments, LysNTA, which contains an extra tail ending in a primary amine group (NH_2_) was used. The complex LysNTA-Ni^2+^ was obtained by mixing 50 mL of a 25 mM solution of LysNTA with 30 mM of NiCl_2_·6H_2_O in a borate-buffered saline (BBS, 50 mM, pH 8.0) during 5 min. The pH was increased with NaOH up to 10.5 to precipitate the excess of Ni^2+^, which was discarded by centrifugation at 5000 xg, RT for 10 min. Then, the pH was adjusted to 9.0, and the complex was stored at 4 °C for future use.

MNPs@PMAO (0.5 mg of Fe) were mixed with LysNTA-Ni^2+^ (32 μmols mg^-1^ Fe) and 18 μmols of α-methoxy-ω-amino polyethylene glycol (PEG, 5000 Da) molecules in a final reaction volume of 1.5 mL. Activation of COOH groups was carried out with 1-ethyl-3-(3-dimethyl aminopropyl)carbodiimide (EDC) in BBS (50 mM, pH 9). The mixture was warmed to 37 °C and 20 μmols of EDC were added to the same reaction tube twice, at time 0 and after 30 min, maintaining the mixture under stirring in a horizontal shaker at 800 rpm for three hours and 30 min. The MNPs were washed with distilled water using centrifugal filters (Amicon, Millipore, 100 kDa cutoff) to eliminate unreacted reagents, then filtered with a Millipore filter (0.22 μm diameter), and stored at 4 °C.

MNPs@PMAO (0.1 mg Fe) previously functionalized with LysNTA-Ni^2+^ and PEG were mixed with E/EC15 fragments (120 or 240 μg mg^-1^ Fe) suspended in a phosphate-buffered saline solution (PBS, w/o Ca^2+^ or Mg^2+^) in a final reaction volume of 150 μL. The bioconjugation was performed by stirring the microtubes on a horizontal shaker for one hour at 37 °C. An extra tube only containing E/EC15 fragments was prepared as the 100 % control to quantify the bioconjugation by Bradford assay following manufacturer’s instructions.

### Physicochemical characterization of MNPs

#### TEM microscopy

The MNP morphology, and size distribution were evaluated by TEM using a Tecnai T20 transmission electron microscope operating at 200 kV. TEM samples were prepared by depositing 5 µl of dilute solution on a copper grid (200 mesh) followed by drying at RT before analysis. MNPs size distributions were obtained by measuring more than 200 MNPs using Fiji software. The concentration used for TEM imaging was 1 µg Fe·mL ¹.

#### Magnetic measurements

The magnetic suspensions were lyophilized and measured as powder, introduced into a gelatine capsule, and immobilized with cotton wool. Hysteresis loops were measured using a superconducting quantum interference device (SQUID, Quantum Design GmbH) magnetometer at 290 K in fields of up to 5000 kA m^-1^.

#### Thermogravimetric analysis

Organic contents on MNPs were determined by thermogravimetric analysis (TGA) using a Universal V4.5A TA Instrument under N_2_ atmosphere at a flow rate of 50 mL min^-1^ at a rate of 10 °C min^-1^ until a final temperature of 800 °C.

#### ICP–AES

The total iron and nickel concentrations were determined. In brief, 20 μL of MNP suspensions were treated with 300 μL of HCl (37 %) solution for 1 h at 80 °C and then diluted with Milli-Q water up to 10 mL and analyzed with a HORIBA Jobin Yvon-ACTIVA-M CCD ICP spectrometer at the SGIker (UPV/EHU) service. Measurements were performed in axial mode with limits of quantification (LOQs) of 0.005 ppm for the two elements. Two wavelengths were used, one to quantify from 0 to 10 ppm and the second to verify that the difference between concentrations using both wavelengths was acceptable with an error of <5 %. The wavelengths were: for iron 238.204 nm (quantitative) and 259.940 nm (confirmation); for nickel 231.604 nm (quantitative) and 216.555 nm (confirmation). Experiments were carried out in duplicate, and results are given as the mean value ± the standard deviation.

#### DLS and ζ-Potential techniques

Measurements were performed on a Malvern Zetasizer Nano instrument considering a refractive index of 2.0 and an absorption index of 1.0 for typical Fe_3_O_4_. Samples were prepared at a concentration of 0.05 mg Fe mL^-1^ and sonicated 5 s before measurement. For the ζ-potential, the dispersed light at an angle of 13° was measured. Each sample was measured at least three times at 25 °C, in water combining at least 5 runs per measurement. Results were treated using the Malvern software Zetasizer Nano 7.13.

### Calcium Aggregation Assay

The bioconjugates (MNPs@E/EC15) were diluted to 100 μg Fe mL^-1^ in PBS (w/o Ca^2+^ or Mg^2+^) or dPBS (0.9 mM Ca^2+^ and Mg^2+^) buffer, and 100 μL were added to a 96-multiwell plate and incubated for 30 min in a horizontal shaker at 37 °C. After this time, the samples were observed under a Nikon ECLIPSE-Ti bright-field microscope. Upon the formation of aggregates in dPBS, EDTA was added reaching a final concentration of 5 mM and the plate was incubated for one hour in a horizontal shaker at 37 °C. The assay was repeated thrice. Images were acquired by using the NIS-Elements software.

### Magnetic field applicator

The magnetic field applicator consisted of a central support for 35 mm Petri dishes and four arrays of magnets located perpendicularly to each other, with each array controlled by an independent servomotor. Neodymium N42 magnets (10 x 10 x 3 mm, Ref. F10103-20), each one with a magnetic field density of 280 mT, were purchased from First4Magnets and used to conform the arrays (five magnets per array). The servomotors were connected to an Arduino Nano board and the corresponding control code was developed in Arduino language. One movement cycle was performed at a frequency of 0.5 Hz. The magnetic flux density was simulated in the software COMSOL Multiphysics.

### Cell culture

MDCK cells were acquired from ATCC (NBL-2; reference: CCL-34). MDCK cells were cultured in DMEM supplemented with 10% (v/v) FBS, GlutaMAX™ (2 mM) and penicillin/streptomycin (100 U mL−1), unless otherwise specified. Cells were maintained at 37°C with 5% CO2 under a humidified atmosphere in sterile conditions.

### Cell labeling with MNPs@E/EC15 bioconjugates and magnetic stimulation

The bioconjugates (MNPs@E/EC15 or MNPs@E/EC12) were diluted to 100 μg Fe mL^-1^ in dPBS (0.9 mM Ca^2+^ and Mg^2+^). Then, the cells were washed once and incubated with PBS for 20 min. Afterwards, 250 μL of the bioconjugate suspensions were added to the cells and incubated for 30 min at 37 °C and 5 % CO_2_ atmosphere. Then, the cells were washed once with dPBS buffer and incubated in DMEM medium or fixed, depending on further steps. For immunofluorescence and SEM experiments, the washed cells were incubated in DMEM medium for 30 min or three hours before fixation.

For immunofluorescence, fixation was performed by adding 200 μL of 2 % paraformaldehyde for 10 min at RT. The cells were then washed three times with PBS, five min per wash.

For SEM imaging, the cells were washed twice with a 0.1 M sodium cacodylate buffer (pH 7.4) and then fixed with a solution of 0.1 M sodium cacodylate and 2 % (v v^-1^) glutaraldehyde at pH 7.4 overnight. Subsequently, the samples were dehydrated by adding 1 mL of methanol solutions of increasing concentration, each for five min, and repeated twice. The percentages of methanol used were 30, 50, 70 and 100 %. Then, the cells were stored in methanol 100 % and sent to the Advanced Microscopy Laboratories (LMA; University of Zaragoza) for coating with palladium (7 nm thickness) and imaging using an INSPECT-F50 SEM microscope. For imaging, 5 and 10 kV were used with the secondary electrons’ detector and the backscatter detector respectively.

For magnetic stimulation, cells were grown in a four-quadrants 35 mm Petri dish (Ibidi, Reference: 80416), labeled with the bioconjugates as described above, and then placed on the magnetic applicator and stimulated for 3 h at 37° C and 5 % CO_2_ atmosphere. Afterwards, cells were incubated for 24 h at 37° C and 5 % CO_2_ atmosphere before lysis for RNA extraction or protein quantification.

### Fluorescence microscopy

All fluorescence microscopy images were obtained on a Nikon ECLIPSE-Ti inverted microscope equipped with a sCMOS camera (Sona 4B, Andor). Appropriate excitation and emission filters were selected for each fluorophore and illumination was performed with a fluorescent lamp (CHGFIE,Nikon).

### Immunofluorescence

Fixed cells were permeabilized by adding 300 μL of a dPBS solution containing 0.25 % (v v^-1^) of Triton-X100 for 10 min. Then, the cells were washed thrice with 200 μL of dPBS, five min per wash. To avoid non-specific labeling, the cells were blocked with a dPBS solution supplemented with 1 % (w v^-1^) of bovine serum albumin (BSA) for 1 h at RT. Then, the cells were washed thrice as before. Subsequently, the cells were incubated with 300 μL of a dPBS solution supplemented with 0.1 % BSA containing a 1:500 dilution of an anti-β-catenin antibody overnight at 4 °C. After this time, the cells were washed thrice and incubated for 40 min with 300 μL of a dPBS solution supplemented with 0.1 % BSA containing a 1:500 dilution of a secondary anti-mouse antibody modified with the fluorophore AlexaFluor 488. Nuclei were stained with 300 μL of a DAPI dilactate solution (1 μg mL^-1^) for 10 min at 37° C, and three final washes were performed with dPBS. Coverslips were mounted using Fluorsave. Images were obtained using the 20X objective.

### High-throughput mRNA extraction, sequencing and data analysis

Total RNA was isolated using Norgen Biotek RNA purification kit. The DNA was removed from total RNAs by digesting in-column with RNAse-free DNAse I according to manufacturer’s instruction. Total RNA quality was evaluated by agarose gel electrophoresis using GelRed for nucleic acid staining. Approximately 1 µg of total RNA per sample (three replicates) were sent to Novogene Bioinformatics Technology for library construction and RNA-sequencing in an Illumina NovaSeq 6000 equipment.

The data alignment was performed using reference genome ensembl_109_canis_lupus_familiaris_ros_cfam_1_0_toplevel. Genes were considered differentially expressed if a Benjamini-Hochberg adjusted P-value was ≤ 0.05 and Log2 fold-change |Log2FC| was ≥ 0.5. Gene ontology and KEGG pathways enrichment analysis were performed using the clusterProfiler package. All computations were performed using R software and the most significant annotations including the presented heatmap were plotted with ggplot2 package.

### MDCK Wnt-luciferase reporter cell line generation

Lentiviral production was performed in HEK-293T cells in a 6 multi-well plate. The corresponding plasmids (pVSV-G, pCMVR8.74 and 7TFP) obtained from Addgene (Items: 138479, 22036, 24308, respectively) were introduced into the cells with Lipofectamine 3000 following manufacturer’s instructions. After 48 h of incubation at 37 °C and 5 % CO_2_ atmosphere, the medium was collected and centrifuged for 10 min at 2000 xg to eliminate cell debris.

A 1:10 dilution of the supernatant was used to ensure an efficient lentiviral transduction of the MDCK cells. The cells were incubated for 6 h with the lentivirus dilution, washed, and incubated with DMEM containing 2.5 µg mL^-1^ of puromycin for 48-72 h. After this time, subcloning was performed by seeding individual cells on a 96 multi-well plate. Briefly, the cells were counted in a Neubauer chamber, and the plates were seeded with a density of 1, 2, 4 and 8 cells per well and incubated until obtaining colonies. Selection of the best clones was carried out by seeding selected colonies on a 24 multi-well plate with a 40 % confluence and incubating for 24 h in DMEM. At this point, Wnt pathway was activated by the addition of Wnt3a ligand (125 ng mL^-1^) diluted in DMEM and the cells were incubated for 24 h. The luciferase activity was measured following the commercial Luciferase Assay System kit (Promega, E1500) and those clones with the highest luciferase activity were selected and frozen in liquid nitrogen for further experiments.

### MDCK Wnt-luciferase activity assay

Luciferase production in the modified MDCK cell line was evaluated from cell lysates, following manufacturer’s instructions. Briefly, MDCK cells were seeded on a four-compartment ibidi with a density of 14 ×10^3^ cells per compartment. After 48 h of incubation in DMEM, at a confluency of 50-60 %, the magnetic stimulation was carried out as described earlier. After 24 h, each compartment containing cells was lysed by adding 150 μL of lysis buffer. In a 96 multi-well plate, 100 μL of luciferase assay reagent and 20 μL of the cell lysate were added and mixed by pipetting twice. The luminescence generated in the enzymatic reaction was measured on a Biotek Synergy H1 UV/vis microplate spectrophotometer. To normalize the luciferase activity, the total protein content on the same lysate was also determined by bicinchoninic acid assay (BCA) following the manufacturer’s instructions. The protein quantification was performed in a lysate dilution (1:5) in water. Luciferase values were normalized by dividing each luminescence measurement by the corresponding protein concentration. The mean value of the control samples was then calculated, and all data (including controls) were normalized to this mean.

### EdU proliferation assay

MDCK cells were seeded on a four-compartment ibidi with a density of 20×10^3^ cells per compartment and incubated in DMEM for 24 h at 37° C and 5% CO_2_ atmosphere. Afterwards, the cells were washed, and the medium was replaced with low glucose-DMEM (1 g L^-1^) without FBS and the culture was incubated in similar conditions for 24 h. Then, the cells were washed once conserving the medium removed (which contained cell signaling molecules whose absence may reduce the proliferation rate),^89^ and were incubated with PBS for 20 min; afterwards, the MNPs@E/EC15 bioconjugates were added to the cells as described earlier and incubated for 30 min. MNPs coated with PMAO without prior modification with TAMRA were used to avoid interference of the fluorescent dye. The cells were washed once with dPBS and the removed medium was mixed with fresh DMEM low glucose (1 g L^-1^) without FBS in a 1:1 ratio adding also the EdU reagent (10 mM). Then, the cells were stimulated with the magnetic applicator for 3 hours and incubated for 15 h at 37° C and 5 % CO_2_ atmosphere. At that time, the cells were fixed and stained following the manufacturer’s instructions. Images were obtained using the 20X objectives.

Five images per condition were processed under the same parameters in ImageJ software to maximize the brightness/contrast ratio. All images were then analyzed using Cell Profiler. The analysis involved identifying nuclei and EdU-positive nuclei as primary objects separately. Then, the percentage of EdU-positive cells was determined by dividing the number of red nuclei (EdU-positive cells) by the number of blue nuclei (total cells) and multiplying by 100.

### Wound healing assay

MDCK cells were seeded on an ibidi containing a two-well culture insert (Ibidi, Reference: 81176) with a density of 17×10^3^ cells per compartment and incubated in DMEM for 48 h at 37° C and 5 % CO_2_ atmosphere. After this time, the cells were washed once with dPBS, the medium was replaced with DMEM low glucose (1 g L^-1^) without FBS, and the culture was incubated in similar conditions for 24 h. Then, the cells were washed once and incubated with PBS for 20 min, the MNPs@E/EC15 were added to the cells as described earlier and incubated for 30 min. The cells were washed once with dPBS and fresh DMEM was added. Then, the cells were stimulated with the magnetic applicator for two hours and incubated for eight hours at 37° C and 5 % CO_2_ atmosphere. Images were obtained in brightfield using the 10X objectives.

Image acquisition was performed every two hours. Cells after magnetic stimulation were taken as the initial time point. Three images were acquired for each independent experiment. Quantification of the area not covered by cells was performed using ImageJ with the *Wound_healing_size_tool* plugin.^90^ Calculations for gap closure percentage and front velocity are detailed in the Calculation S1 (Supporting Information).

### Statistical analysis

Data were analyzed using GraphPad Prism 9.0 (GraphPad Software, San Diego, USA). Each independent experiment corresponds to an experiment performed in different days. In all cases, at least two independent experiments performed in different days were evaluated. Analysis of variance (ANOVA) one-way with Tukey’s comparison test were used to evaluate differences between groups, and differences were considered statistically significant at p-value<0.05. Simple t-test was also used for comparisons between two groups, and differences were considered statistically significant at p-value<0.05.

## Supporting Information

The Supporting Information is available free of charge and consists of:

Additional characterization of protein fragments, functionalization of MNPs, and bioconjugates as well as images of the cell labeling with bioconjugates containing either E/EC12 or E/EC15 fragments. A theoretical simulation of the magnetic field in the center of the cell culture dish during magnets movement. Immunofluorescence of MDCK cells labeled with E-cadherin bioconjugates and stimulated with magnets. RNA-seq additional information regarding intra and intergroup variability, additional categories of GO and KEGG enrichment analysis, Heatmap of DEGs from ‘MNPs + Magnet’ vs MNPs comparison, flow cytometry analysis of E-cadherin expression, and images of EdU staining as well as the method and primer sequences used for RT-qPCR are also available.

## Acknowledgements

The authors would like to acknowledge the use of Servicios Cientifícos Técnicos del CIBA (IACS-Universidad de Zaragoza), the Advanced Microscopy Laboratory (Universidad de Zaragoza), for access to their instrumentation and expertise and for the use of Servicio General de Apoyo a la Investigación-SAI, Universidad de Zaragoza. This work has been supported by Grant CEX2023-001286-S funded by MICIU/AEI /10.13039/501100011033. L.G-R acknowledges Gobierno de Aragón for predoctoral fellowship (2024–2028). The authors also acknowledge support from Gobierno de Aragón and “ERDF A way of making Europe” for funding the Bionanosurf (E15_20R) research group and the use of Servicio General de Apoyo a la Investigación-SAI, Universidad de Zaragoza. We sincerely thanks Dr. Daniel García Gonzalez (Universidad Carlos III, Madrid) for generously providing the magnetic stimulator prototype which was instrumental in the final design of our experimental setup.

## Conflict of Interest

The authors declare no conflict of interest

## Data Availability Statement

The transcriptomic data that support the findings of this study are available at ArrayExpress with the accession number: E-MTAB-15853; or from the corresponding author upon reasonable request.

## Funding Sources

This work has received funding from the European Research Council (ERC) under the European Union’s Horizon 2020 research and innovation programme (Grant agreement No. 853468) and Agencia Estatal de Investigación, Project PID2021-122508NB-I00 funded by MICIU/AEI/10.13039/501100011033 and FEDER, and from Gobierno de Aragón (funding for the Bionanosurf research group E15_23R).

## Notes

### Competing Interest Statement

The authors have declared no competing interest.

## References

(1) Spencer, D. M.; Wandless, T. J.; Schreiber, S. L.; Crabtree, G. R. Controlling Signal Transduction with Synthetic Ligands. Science 1993, 262 (5136), 1019–1024. 10.1126/science.7694365.

(2) Handly, L. N.; Yao, J.; Wollman, R. Signal Transduction at the Single-Cell Level: Approaches to Study the Dynamic Nature of Signaling Networks. J Mol Biol 2016, 428 (19), 3669–3682. 10.1016/j.jmb.2016.07.009.

(3) Shin, W.; Jeong, S.; Lee, J.; Jeong, S. Y.; Shin, J.; Kim, H. H.; Cheon, J.; Lee, J.-H. Magnetogenetics with Piezo1 Mechanosensitive Ion Channel for CRISPR Gene Editing. Nano Lett 2022, 22 (18), 7415–7422. 10.1021/acs.nanolett.2c02314.

(4) Di, X.; Gao, X.; Peng, L.; Ai, J.; Jin, X.; Qi, S.; Li, H.; Wang, K.; Luo, D. Cellular Mechanotransduction in Health and Diseases: From Molecular Mechanism to Therapeutic Targets. Signal Transduct Target Ther 2023, 8 (1), 282. 10.1038/s41392-023-01501-9.

(5) Cao, R.; Tian, H.; Tian, Y.; Fu, X. A Hierarchical Mechanotransduction System: From Macro to Micro. Advanced Science 2024, 11 (11). 10.1002/advs.202302327.

(6) Del Sol-Fernández, S.; Martínez-Vicente, P.; Gomollón-Zueco, P.; Castro-Hinojosa, C.; Gutiérrez, L.; Fratila, R. M.; Moros, M. Magnetogenetics: Remote Activation of Cellular Functions Triggered by Magnetic Switches. Nanoscale 2022, 14 (6), 2091–2118. 10.1039/d1nr06303k.

(7) Liu, A. P. Biophysical Tools for Cellular and Subcellular Mechanical Actuation of Cell Signaling. Biophysical Journal. Biophysical Society September 20, 2016, pp 1112–1118. 10.1016/j.bpj.2016.02.043.

(8) Bielfeldt, M.; Rebl, H.; Peters, K.; Sridharan, K.; Staehlke, S.; Nebe, J. B. Sensing of Physical Factors by Cells: Electric Field, Mechanical Forces, Physical Plasma and Light—Importance for Tissue Regeneration. Biomedical Materials and Devices. Springer Nature March 1, 2023, pp 146–161. 10.1007/s44174-022-00028-x.

(9) Jin, P.; Jan, L. Y.; Jan, Y.-N. Mechanosensitive Ion Channels: Structural Features Relevant to Mechanotransduction Mechanisms. Annu Rev Neurosci 2020, 43 (1), 207–229. 10.1146/annurev-neuro-070918-050509.

(10) Roeterink, R. M. A.; Casadevall i Solvas, X.; Collins, D. J.; Scott, D. J. Force versus Response: Methods for Activating and Characterizing Mechanosensitive Ion Channels and GPCRs. Adv Healthc Mater 2024, 13 (31). 10.1002/adhm.202402167.

(11) Seo, D.; Southard, K. M.; Kim, J.; Lee, H. J.; Farlow, J.; Lee, J.; Litt, D. B.; Haas, T.; Alivisatos, A. P.; Cheon, J.; Gartner, Z. J.; Jun, Y. A Mechanogenetic Toolkit for Interrogating Cell Signaling in Space and Time. Cell 2016, 165 (6), 1507–1518. 10.1016/j.cell.2016.04.045.

(12) Teo, J.-L.; Kahn, M. The Wnt Signaling Pathway in Cellular Proliferation and Differentiation: A Tale of Two Coactivators. Adv Drug Deliv Rev 2010, 62 (12), 1149–1155. 10.1016/j.addr.2010.09.012.

(13) Houschyar, K. S.; Momeni, A.; Pyles, M. N.; Maan, Z. N.; Whittam, A. J.; Siemers, F. Wnt Signaling Induces Epithelial Differentiation during Cutaneous Wound Healing. Organogenesis 2015, 11 (3), 95–104. 10.1080/15476278.2015.1086052.

(14) MacDonald, B. T.; Tamai, K.; He, X. Wnt/β-Catenin Signaling: Components, Mechanisms, and Diseases. Dev Cell 2009, 17 (1), 9–26. 10.1016/j.devcel.2009.06.016.

(15) Liu, J.; Xiao, Q.; Xiao, J.; Niu, C.; Li, Y.; Zhang, X.; Zhou, Z.; Shu, G.; Yin, G. Wnt/β-Catenin Signalling: Function, Biological Mechanisms, and Therapeutic Opportunities. Signal Transduct Target Ther 2022, 7 (1), 3. 10.1038/s41392-021-00762-6.

(16) DiRenzo, D. M.; Chaudhary, M. A.; Shi, X.; Franco, S. R.; Zent, J.; Wang, K.; Guo, L.-W.; Kent, K. C. A Crosstalk between TGF-β/Smad3 and Wnt/β-Catenin Pathways Promotes Vascular Smooth Muscle Cell Proliferation. Cell Signal 2016, 28 (5), 498–505. 10.1016/j.cellsig.2016.02.011.

(17) Shapiro, M.; Akiri, G.; Chin, C.; Wisnivesky, J. P.; Beasley, M. B.; Weiser, T. S.; Swanson, S. J.; Aaronson, S. A. Wnt Pathway Activation Predicts Increased Risk of Tumor Recurrence in Patients With Stage I Nonsmall Cell Lung Cancer. Ann Surg 2013, 257 (3), 548–554. 10.1097/SLA.0b013e31826d81fd.

(18) Zhan, T.; Rindtorff, N.; Boutros, M. Wnt Signaling in Cancer. Oncogene 2017, 36 (11), 1461–1473. 10.1038/onc.2016.304.

(19) He, S.; Lu, Y.; Liu, X.; Huang, X.; Keller, E. T.; Qian, C.-N.; Zhang, J. Wnt3a: Functions and Implications in Cancer. Chin J Cancer 2015, 34 (3), 50. 10.1186/s40880-015-0052-4.

(20) Huang, P.; Yan, R.; Zhang, X.; Wang, L.; Ke, X.; Qu, Y. Activating Wnt/β-Catenin Signaling Pathway for Disease Therapy: Challenges and Opportunities. Pharmacol Ther 2019, 196, 79–90. 10.1016/j.pharmthera.2018.11.008.

(21) Bonnet, C.; Brahmbhatt, A.; Deng, S. X.; Zheng, J. J. Wnt Signaling Activation: Targets and Therapeutic Opportunities for Stem Cell Therapy and Regenerative Medicine. RSC Chem Biol 2021, 2 (4), 1144–1157. 10.1039/D1CB00063B.

(22) Repina, N. A.; Johnson, H. J.; Bao, X.; Zimmermann, J. A.; Joy, D. A.; Bi, S. Z.; Kane, R. S.; Schaffer, D. V. Optogenetic Control of Wnt Signaling Models Cell-Intrinsic Embryogenic Patterning Using 2D Human Pluripotent Stem Cell Culture. Development 2023, 150 (14). 10.1242/dev.201386.

(23) Liu, D.-X.; Hao, S.-L.; Yang, W.-X. Crosstalk Between β-CATENIN-Mediated Cell Adhesion and the Wnt Signaling Pathway. DNA Cell Biol 2023, 42 (1), 1–13. 10.1089/dna.2022.0424.

(24) Yulis, M.; Kusters, D. H. M.; Nusrat, A. Cadherins: Cellular Adhesive Molecules Serving as Signalling Mediators. J Physiol 2018, 596 (17), 3883–3898. 10.1113/JP275328.

(25) Leckband, D. E.; de Rooij, J. Cadherin Adhesion and Mechanotransduction. Annu Rev Cell Dev Biol 2014, 30 (1), 291–315. 10.1146/annurev-cellbio-100913-013212.

(26) Yap, A.; Liang, X.; Gomez, G. Current Perspectives on Cadherin-Cytoskeleton Interactions and Dynamics. Cell Health Cytoskelet 2015, 7, 11. 10.2147/CHC.S76107.

(27) Röper, J.-C.; Mitrossilis, D.; Stirnemann, G.; Waharte, F.; Brito, I.; Fernandez-Sanchez, M.-E.; Baaden, M.; Salamero, J.; Farge, E. The Major β-Catenin/E-Cadherin Junctional Binding Site Is a Primary Molecular Mechano-Transductor of Differentiation *in vivo*. Elife 2018, 7. 10.7554/eLife.33381.

(28) Benham-Pyle, B. W.; Pruitt, B. L.; Nelson, W. J. Mechanical Strain Induces E-Cadherin–Dependent Yap1 and β-Catenin Activation to Drive Cell Cycle Entry. Science 2015, 348 (6238), 1024–1027. 10.1126/science.aaa4559.

(29) Benham-Pyle, B. W.; Sim, J. Y.; Hart, K. C.; Pruitt, B. L.; Nelson, W. J. Increasing β-Catenin/Wnt3A Activity Levels Drive Mechanical Strain-Induced Cell Cycle Progression through Mitosis. Elife 2016, 5. 10.7554/eLife.19799.

(30) Stockinger, A.; Eger, A.; Wolf, J.; Beug, H.; Foisner, R. E-Cadherin Regulates Cell Growth by Modulating Proliferation-Dependent β-Catenin Transcriptional Activity. Journal of Cell Biology 2001, 154 (6), 1185–1196. 10.1083/jcb.200104036.

(31) Pinheiro, D.; Bellaïche, Y. Mechanical Force-Driven Adherens Junction Remodeling and Epithelial Dynamics. Dev Cell 2018, 47 (1), 3–19. 10.1016/j.devcel.2018.09.014.

(32) Viji Babu, P. K.; Mirastschijski, U.; Belge, G.; Radmacher, M. Homophilic and Heterophilic Cadherin Bond Rupture Forces in Homo- or Hetero-Cellular Systems Measured by AFM-Based Single-Cell Force Spectroscopy. European Biophysics Journal 2021, 50 (3–4), 543–559. 10.1007/s00249-021-01536-2.

(33) Muhamed, I.; Wu, J.; Sehgal, P.; Kong, X.; Tajik, A.; Wang, N.; Leckband, D. E. E-Cadherin-Mediated Force Transduction Signals Regulate Global Cell Mechanics. J Cell Sci 2016, 129 (9), 1843–1854. 10.1242/jcs.185447.

(34) le Duc, Q.; Shi, Q.; Blonk, I.; Sonnenberg, A.; Wang, N.; Leckband, D.; de Rooij, J. Vinculin Potentiates E-Cadherin Mechanosensing and Is Recruited to Actin-Anchored Sites within Adherens Junctions in a Myosin II–Dependent Manner. Journal of Cell Biology 2010, 189 (7), 1107–1115. 10.1083/jcb.201001149.

(35) Bays, J. L.; Campbell, H. K.; Heidema, C.; Sebbagh, M.; DeMali, K. A. Linking E-Cadherin Mechanotransduction to Cell Metabolism through Force-Mediated Activation of AMPK. Nat Cell Biol 2017, 19 (6), 724–731. 10.1038/ncb3537.

(36) Jasaitis, A.; Estevez, M.; Heysch, J.; Ladoux, B.; Dufour, S. E-Cadherin-Dependent Stimulation of Traction Force at Focal Adhesions via the Src and PI3K Signaling Pathways. Biophys J 2012, 103 (2), 175–184. 10.1016/j.bpj.2012.06.009.

(37) Perez, T. D.; Tamada, M.; Sheetz, M. P.; Nelson, W. J. Immediate-Early Signaling Induced by E-Cadherin Engagement and Adhesion. Journal of Biological Chemistry 2008, 283 (8), 5014–5022. 10.1074/jbc.M705209200.

(38) Sehgal, P.; Kong, X.; Wu, J.; Sunyer, R.; Trepat, X.; Leckband, D. Epidermal Growth Factor Receptor and Integrins Control Force-Dependent Vinculin Recruitment to E-Cadherin Junctions. J Cell Sci 2018, 131 (6). 10.1242/jcs.206656.

(39) Qin, E. C.; Ahmed, S. T.; Sehgal, P.; Vu, V. H.; Kong, H.; Leckband, D. E. Comparative Effects of N-Cadherin Protein and Peptide Fragments on Mesenchymal Stem Cell Mechanotransduction and Paracrine Function. Biomaterials 2020, 239, 119846. 10.1016/j.biomaterials.2020.119846.

(40) Castro-Hinojosa, C.; Del Sol-Fernández, S.; Moreno-Antolín, E.; Martín-Gracia, B.; Ovejero, J. G.; de la Fuente, J. M.; Grazú, V.; Fratila, R. M.; Moros, M. A Simple and Versatile Strategy for Oriented Immobilization of His-Tagged Proteins on Magnetic Nanoparticles. Bioconjug Chem 2023, 34 (12), 2275–2292. 10.1021/acs.bioconjchem.3c00417.

(41) Harrison, O. J.; Corps, E. M.; Kilshaw, P. J. Cadherin Adhesion Depends on a Salt Bridge at the N-Terminus. J Cell Sci 2005, 118 (18), 4123–4130. 10.1242/jcs.02539.

(42) Vendome, J.; Felsovalyi, K.; Song, H.; Yang, Z.; Jin, X.; Brasch, J.; Harrison, O. J.; Ahlsen, G.; Bahna, F.; Kaczynska, A.; Katsamba, P. S.; Edmond, D.; Hubbell, W. L.; Shapiro, L.; Honig, B. Structural and Energetic Determinants of Adhesive Binding Specificity in Type I Cadherins. Proceedings of the National Academy of Sciences 2014, 111 (40), E4175–E4184. 10.1073/pnas.1416737111.

(43) Zhao, H.; Liang, Y.; Xu, Z.; Wang, L.; Zhou, F.; Li, Z.; Jin, J.; Yang, Y.; Fang, Z.; Hu, Y.; Zhang, L.; Su, J.; Zha, X. N Glycosylation Affects the Adhesive Function of E Cadherin through Modifying the Composition of Adherens Junctions (AJs) in Human Breast Carcinoma Cell Line MDA MB 435. J Cell Biochem 2008, 104 (1), 162–175. 10.1002/jcb.21608.

(44) Perret, E.; Leung, A.; Feracci, H.; Evans, E. Trans-Bonded Pairs of E-Cadherin Exhibit a Remarkable Hierarchy of Mechanical Strengths; 2004; Vol. 101. www.pnas.orgcgidoi10.1073pnas.0402085101.

(45) Wingfield, P. T. Overview of the Purification of Recombinant Proteins. Curr Protoc Protein Sci 2015, 80 (1), 6.1.1–6.1.35. 10.1002/0471140864.ps0601s80.

(46) Perkins, D. N.; Pappin, D. J. C.; Creasy, D. M.; Cottrell, J. S. Probability-Based Protein Identification by Searching Sequence Databases Using Mass Spectrometry Data. Electrophoresis 1999, 20 (18), 3551–3567. https://doi.org/10.1002/(SICI)1522-2683(19991201)20:18<3551::AID-ELPS3551>3.0.CO;2-2.

(47) Stone, K. L.; Deangelis, R.; LoPresti, M.; Jones, J.; Papov, V. V.; Williams, K. R. Use of Liquid Chromatography electrospray Ionization tandem Mass Spectrometry (LC ESI MS/MS) for Routine Identification of Enzymatically Digested Proteins Separated by Sodium Dodecyl Sulfate polyacrylamide Gel Electrophoresis. Electrophoresis 1998, 19 (6), 1046–1052. 10.1002/elps.1150190620.

(48) Shapiro, L.; Weis, W. I. Structure and Biochemistry of Cadherins and Catenins. Cold Spring Harb Perspect Biol 2009, 1 (3), a003053–a003053. 10.1101/cshperspect.a003053.

(49) Del Sol Fernández, S.; De Simone, M.; Fernández Afonso, Y.; Garcia Gonzalez, D.; Martínez Vicente, P.; van Zanten, T.; Fratila, R. M.; Moros, M. MagPiezo: A Magnetogenetic Platform for Remote Activation of Endogenous Piezo1 Channels in Endothelial Cells. Adv Funct Mater 2026. 10.1002/adfm.202529076.

(50) Moros, M.; Hernáez, B.; Garet, E.; Dias, J. T.; Sáez, B.; Grazú, V.; González-Fernández, Á.; Alonso, C.; de la Fuente, J. M. Monosaccharides versus PEG-Functionalized NPs: Influence in the Cellular Uptake. ACS Nano 2012, 6 (2), 1565–1577. 10.1021/nn204543c.

(51) Moros, M.; Pelaz, B.; López-Larrubia, P.; García-Martin, M. L.; Grazú, V.; de la Fuente, J. M. Engineering Biofunctional Magnetic Nanoparticles for Biotechnological Applications. Nanoscale 2010, 2 (9), 1746. 10.1039/c0nr00104j.

(52) Tomasello, G.; Armenia, I.; Molla, G. The Protein Imager: A Full-Featured Online Molecular Viewer Interface with Server-Side HQ-Rendering Capabilities. Bioinformatics 2020, 36 (9), 2909–2911. 10.1093/bioinformatics/btaa009.

(53) Masthoff, I.-C.; David, F.; Wittmann, C.; Garnweitner, G. Functionalization of Magnetic Nanoparticles with High-Binding Capacity for Affinity Separation of Therapeutic Proteins. Journal of Nanoparticle Research 2014, 16 (1), 2164. 10.1007/s11051-013-2164-6.

(54) Kim, J. S.; Valencia, C. A.; Liu, R.; Lin, W. Highly-Efficient Purification of Native Polyhistidine-Tagged Proteins by Multivalent NTA-Modified Magnetic Nanoparticles. Bioconjug Chem 2007, 18 (2), 333–341. 10.1021/bc060195l.

(55) Emond, M. R.; Jontes, J. D. Bead Aggregation Assays for the Characterization of Putative Cell Adhesion Molecules. Journal of Visualized Experiments 2014, No. 92, 1–6. 10.3791/51762.

(56) Chappuis-Flament, S.; Wong, E.; Hicks, L. D.; Kay, C. M.; Gumbiner, B. M. Multiple Cadherin Extracellular Repeats Mediate Homophilic Binding and Adhesion. J Cell Biol 2001, 154 (1), 231–243. 10.1083/jcb.200103143.

(57) Chen, Y.-T.; Stewart, D. B.; Nelson, W. J. Coupling Assembly of the E-Cadherin/β-Catenin Complex to Efficient Endoplasmic Reticulum Exit and Basal-Lateral Membrane Targeting of E-Cadherin in Polarized MDCK Cells. J Cell Biol 1999, 144 (4), 687–699. 10.1083/jcb.144.4.687.

(58) Moreno-Mateos, M. A.; Gonzalez-Rico, J.; Nunez-Sardinha, E.; Gomez-Cruz, C.; Lopez-Donaire, M. L.; Lucarini, S.; Arias, A.; Muñoz-Barrutia, A.; Velasco, D.; Garcia-Gonzalez, D. Magneto-Mechanical System to Reproduce and Quantify Complex Strain Patterns in Biological Materials. Appl Mater Today 2022, 27, 101437. 10.1016/j.apmt.2022.101437.

(59) Maldonado-Camargo, L.; Unni, M.; Rinaldi, C. Magnetic Characterization of Iron Oxide Nanoparticles for Biomedical Applications. In Methods in Molecular Biology; Humana Press Inc., 2017; Vol. 1570, pp 47–71. 10.1007/978-1-4939-6840-4_4.

(60) Ansari, K.; Ahmad, R.; Tanweer, M. S.; Azam, I. Magnetic Iron Oxide Nanoparticles as a Tool for the Advancement of Biomedical and Environmental Application: A Review. Biomedical Materials & Devices 2024, 2 (1), 139–157. 10.1007/s44174-023-00091-y.

(61) Kahmann, T.; Ludwig, F. Magnetic Field Dependence of the Effective Magnetic Moment of Multi-Core Nanoparticles. J Appl Phys 2020, 127 (23). 10.1063/5.0011629.

(62) Carrey, J.; Hallali, N. Torque Undergone by Assemblies of Single-Domain Magnetic Nanoparticles Submitted to a Rotating Magnetic Field. Phys Rev B 2016, 94 (18), 184420. 10.1103/PhysRevB.94.184420.

(63) Shen, Y.; Wu, C.; Uyeda, T. Q. P.; Plaza, G. R.; Liu, B.; Han, Y.; Lesniak, M. S.; Cheng, Y. Elongated Nanoparticle Aggregates in Cancer Cells for Mechanical Destruction with Low Frequency Rotating Magnetic Field. Theranostics 2017, 7 (6), 1735–1748. 10.7150/thno.18352.

(64) Wu, J.; Goyal, R.; Grandl, J. Localized Force Application Reveals Mechanically Sensitive Domains of Piezo1. Nat Commun 2016, 7 (1), 12939. 10.1038/ncomms12939.

(65) Lee, J.; Shin, W.; Lim, Y.; Kim, J.; Kim, W. R.; Kim, H.; Lee, J.-H.; Cheon, J. Non-Contact Long-Range Magnetic Stimulation of Mechanosensitive Ion Channels in Freely Moving Animals. Nat Mater 2021, 20 (7), 1029–1036. 10.1038/s41563-020-00896-y.

(66) Borghi, N.; Sorokina, M.; Shcherbakova, O. G.; Weis, W. I.; Pruitt, B. L.; Nelson, W. J.; Dunn, A. R. E-Cadherin Is under Constitutive Actomyosin-Generated Tension That Is Increased at Cell–Cell Contacts upon Externally Applied Stretch. Proceedings of the National Academy of Sciences 2012, 109 (31), 12568–12573. 10.1073/pnas.1204390109.

(67) Koch, C. M.; Chiu, S. F.; Akbarpour, M.; Bharat, A.; Ridge, K. M.; Bartom, E. T.; Winter, D. R. A Beginner’s Guide to Analysis of RNA Sequencing Data. Am J Respir Cell Mol Biol 2018, 59 (2), 145–157. 10.1165/rcmb.2017-0430TR.

(68) Ashburner, M.; Ball, C. A.; Blake, J. A.; Botstein, D.; Butler, H.; Cherry, J. M.; Davis, A. P.; Dolinski, K.; Dwight, S. S.; Eppig, J. T.; Harris, M. A.; Hill, D. P.; Issel-Tarver, L.; Kasarskis, A.; Lewis, S.; Matese, J. C.; Richardson, J. E.; Ringwald, M.; Rubin, G. M.; Sherlock, G. Gene Ontology: Tool for the Unification of Biology. Nat Genet 2000, 25 (1), 25–29. 10.1038/75556.

(69) Buckley, C. D.; Tan, J.; Anderson, K. L.; Hanein, D.; Volkmann, N.; Weis, W. I.; Nelson, W. J.; Dunn, A. R. The Minimal Cadherin-Catenin Complex Binds to Actin Filaments under Force. Science (1979) 2014, 346 (6209). 10.1126/science.1254211.

(70) Kannan, N.; Tang, V. W. Myosin-1c Promotes E-Cadherin Tension and Force-Dependent Recruitment of α-Actinin to the Epithelial Cell Junction. J Cell Sci 2018, 131 (12). 10.1242/jcs.211334.

(71) Hultin, S.; Subramani, A.; Hildebrand, S.; Zheng, Y.; Majumdar, A.; Holmgren, L. AmotL2 Integrates Polarity and Junctional Cues to Modulate Cell Shape. Sci Rep 2017, 7 (1), 7548. 10.1038/s41598-017-07968-1.

(72) Verma, S.; Han, S. P.; Michael, M.; Gomez, G. A.; Yang, Z.; Teasdale, R. D.; Ratheesh, A.; Kovacs, E. M.; Ali, R. G.; Yap, A. S. A WAVE2–Arp2/3 Actin Nucleator Apparatus Supports Junctional Tension at the Epithelial Zonula Adherens. Mol Biol Cell 2012, 23 (23), 4601–4610. 10.1091/mbc.e12-08-0574.

(73) Kehrberg, R. J.; DeMali, K. A. E-Cadherin: A Conductor of Cellular Signaling. Curr Opin Cell Biol 2025, 95, 102559. 10.1016/j.ceb.2025.102559.

(74) Xue, C.; Chu, Q.; Shi, Q.; Zeng, Y.; Lu, J.; Li, L. Wnt Signaling Pathways in Biology and Disease: Mechanisms and Therapeutic Advances. Signal Transduct Target Ther 2025, 10 (1), 106. 10.1038/s41392-025-02142-w.

(75) Basson, M. A. Signaling in Cell Differentiation and Morphogenesis. Cold Spring Harb Perspect Biol 2012, 4 (6), a008151–a008151. 10.1101/cshperspect.a008151.

(76) Deng, Z.; Fan, T.; Xiao, C.; Tian, H.; Zheng, Y.; Li, C.; He, J. TGF-β Signaling in Health, Disease and Therapeutics. Signal Transduct Target Ther 2024, 9 (1), 61. 10.1038/s41392-024-01764-w.

(77) Izaguirre, M. F.; Larrea, D.; Adur, J. F.; Diaz-Zamboni, J. E.; Vicente, N. B.; Galetto, C. D.; Casco, V. H. Role of E-Cadherin in Epithelial Architecture Maintenance. Cell Commun Adhes 2010, 17 (1), 1–12. 10.3109/15419061003686938.

(78) Abuammah, A.; Maimari, N.; Towhidi, L.; Frueh, J.; Chooi, K. Y.; Warboys, C.; Krams, R. New Developments in Mechanotransduction: Cross Talk of the Wnt, TGF-β and Notch Signalling Pathways in Reaction to Shear Stress. Curr Opin Biomed Eng 2018, 5, 96–104. 10.1016/j.cobme.2018.03.003.

(79) Luo, K. Signaling Cross Talk between TGF-β/Smad and Other Signaling Pathways. Cold Spring Harb Perspect Biol 2017, 9 (1), a022137. 10.1101/cshperspect.a022137.

(80) Moshkovsky, A. R.; Kirschner, M. W. The Nonredundant Nature of the Axin2 Regulatory Network in the Canonical Wnt Signaling Pathway. Proceedings of the National Academy of Sciences 2022, 119 (9). 10.1073/pnas.2108408119.

(81) Röhrs, S.; Kutzner, N.; Vlad, A.; Grunwald, T.; Ziegler, S.; Müller, O. Chronological Expression of Wnt Target Genes *Ccnd1*, *Myc*, *Cdkn1a*, *Tfrc*, *Plf1*, and *Ramp3*. Cell Biol Int 2009, 33 (4), 501–508. 10.1016/j.cellbi.2009.01.016.

(82) Fuerer, C.; Nusse, R. Lentiviral Vectors to Probe and Manipulate the Wnt Signaling Pathway. PLoS One 2010, 5 (2), e9370. 10.1371/journal.pone.0009370.

(83) Cai, D.; Chen, S.-C.; Prasad, M.; He, L.; Wang, X.; Choesmel-Cadamuro, V.; Sawyer, J. K.; Danuser, G.; Montell, D. J. Mechanical Feedback through E-Cadherin Promotes Direction Sensing during Collective Cell Migration. Cell 2014, 157 (5), 1146–1159. 10.1016/j.cell.2014.03.045.

(84) Rotherham, M.; Nahar, T.; Goodman, T.; Telling, N.; Gates, M.; El Haj, A. Magnetic Mechanoactivation of Wnt Signaling Augments Dopaminergic Differentiation of Neuronal Cells. Adv Biosyst 2019, 3 (9). 10.1002/adbi.201900091.

(85) Rotherham, M.; El Haj, A. J. Remote Activation of the Wnt/β-Catenin Signalling Pathway Using Functionalised Magnetic Particles. PLoS One 2015, 10 (3), e0121761. 10.1371/journal.pone.0121761.

(86) Hu, B.; Rotherham, M.; Farrow, N.; Roach, P.; Dobson, J.; El Haj, A. J. Immobilization of Wnt Fragment Peptides on Magnetic Nanoparticles or Synthetic Surfaces Regulate Wnt Signaling Kinetics. Int J Mol Sci 2022, 23 (17), 10164. 10.3390/ijms231710164.

(87) Dias, J. T.; Moros, M.; del Pino, P.; Rivera, S.; Grazú, V.; de la Fuente, J. M. DNA as a Molecular Local Thermal Probe for the Analysis of Magnetic Hyperthermia. Angewandte Chemie International Edition 2013, 52 (44), 11526–11529. 10.1002/anie.201305835.

(88) Yoe, J. H.; Jones, A. LETCHER. Colorimetric Determination of Iron with Disodium-1,2-Dihydroxybenzene-3,5-Disulfonate. Industrial & Engineering Chemistry Analytical Edition 1944, 16 (2), 111–115. 10.1021/i560126a015.

(89) Su, J.; Song, Y.; Zhu, Z.; Huang, X.; Fan, J.; Qiao, J.; Mao, F. Cell–Cell Communication: New Insights and Clinical Implications. Signal Transduct Target Ther 2024, 9 (1), 196. 10.1038/s41392-024-01888-z.

(90) Suarez-Arnedo, A.; Torres Figueroa, F.; Clavijo, C.; Arbeláez, P.; Cruz, J. C.; Muñoz-Camargo, C. An Image J Plugin for the High Throughput Image Analysis of in Vitro Scratch Wound Healing Assays. PLoS One 2020, 15 (7), e0232565. 10.1371/journal.pone.0232565.

