## Supporting information for "Remote Activation of Wnt Signaling and Cell Proliferation by E-cadherin Magnetomechanical Stimulation"

Index

### 1. **E/EC15 cadherin fragments identification**

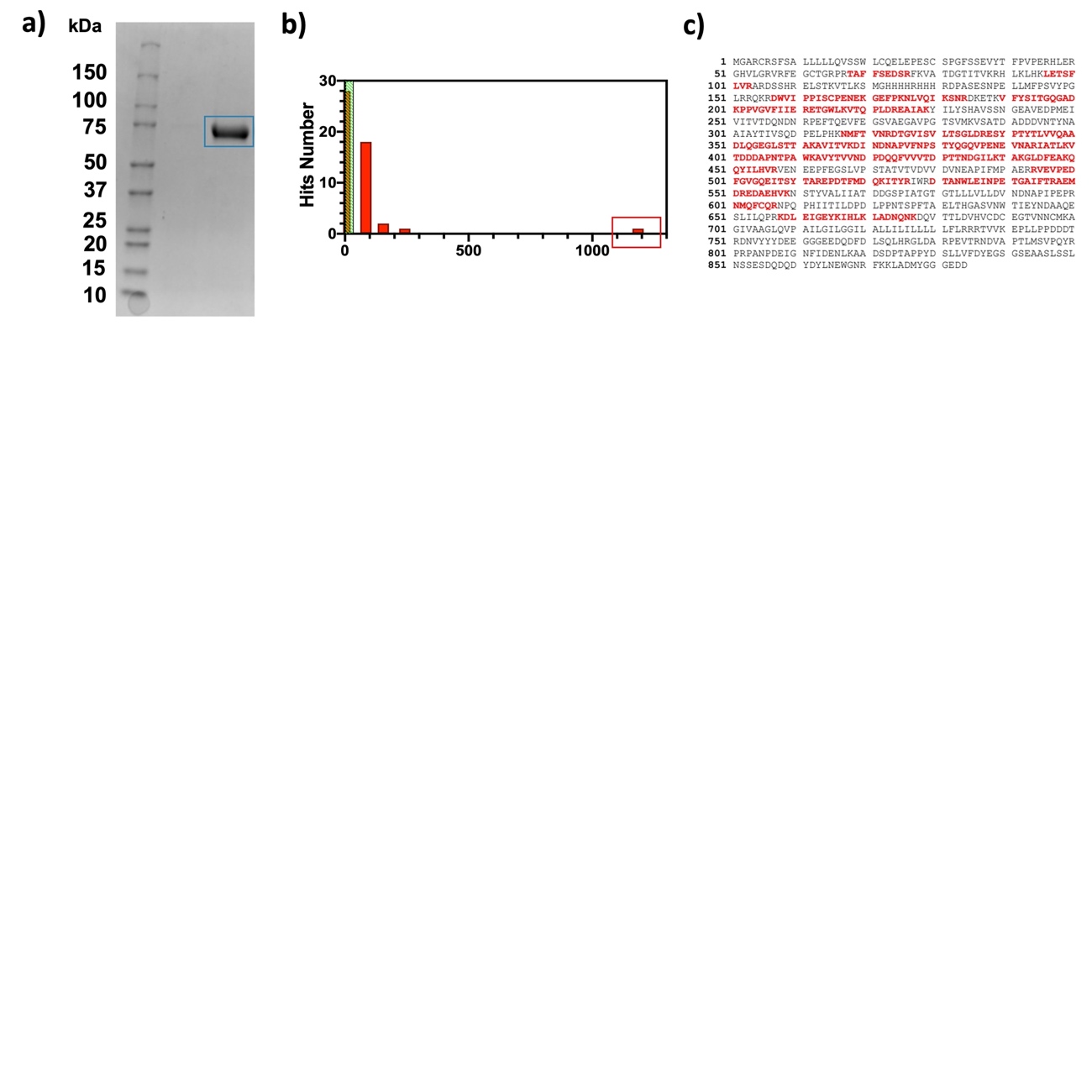

**Figure S1.** a) SDS-PAGE showing protein isolated using IMAC for His-tagged proteins in a chromatography system. b) Mascot protein score. The highest score (1185) corresponds to E-cadherin (red box). c) Sequence coverage map of E-cadherin highlighting in red the peptides identified by liquid chromatography-electrospray ionization-mass spectrometry (LC-ESI-MS) after trypsin digestion of the protein band excised from the SDS-PAGE gel.

### 2. **MNP size determination**

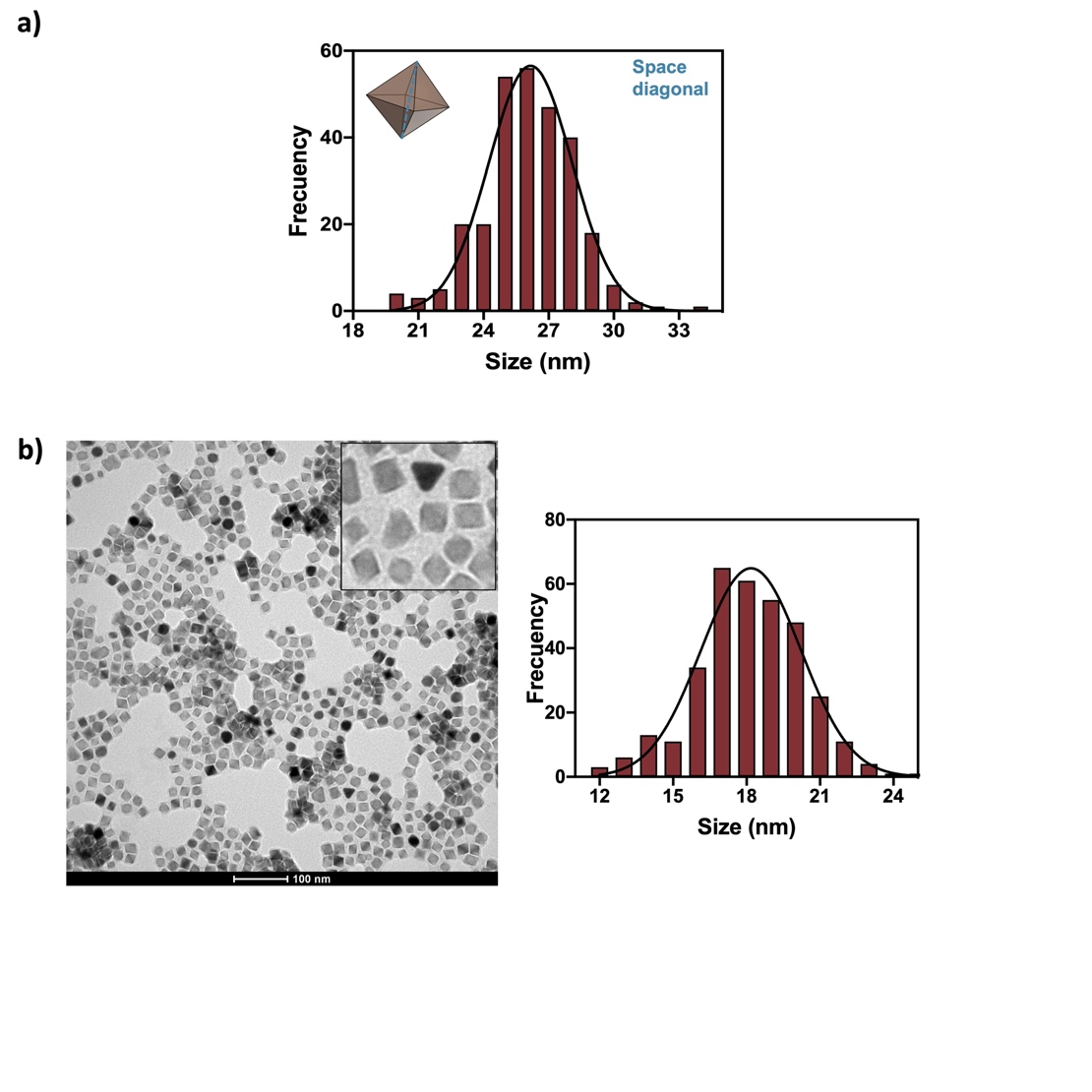

**Figure S2.** a) Space diagonal size distribution of MNPs in hexane, determined by measuring 350 particles and fitting to a Gaussian curve, the data was obtained from the image showed in Figure 2b. b) TEM image of MNPs@PMAO and their edge size distribution by measuring 300 particles and fitting to a Gaussian curve; mean size: 18.2 ± 2.0 nm; Scale bar: 100 nm.

### 3. **Functionalization of MNPs with PEG + LysNTA-Ni^2+^**

**Table S1.** Hydrodynamic diameter by number and ζ-potential after each functionalization step

| Functionalization step | Diameter by number  (nm) | ζ -potential  (mV) |
| --- | --- | --- |
| MNPs@PMAO | 78 ± 10 | -25 ± 1 |
| MNPs@PMAO  @(PEG5000 + LysNTA-Ni^2+^) | 124 ± 18 | -16 ± 1 |
| MNPs@E/EC15 | 114 ± 21 | -14 ± 1 |

### 4. **MDCK cell labeling with MNPs@E/EC12 and MNPs@E/EC15 bioconjugates**

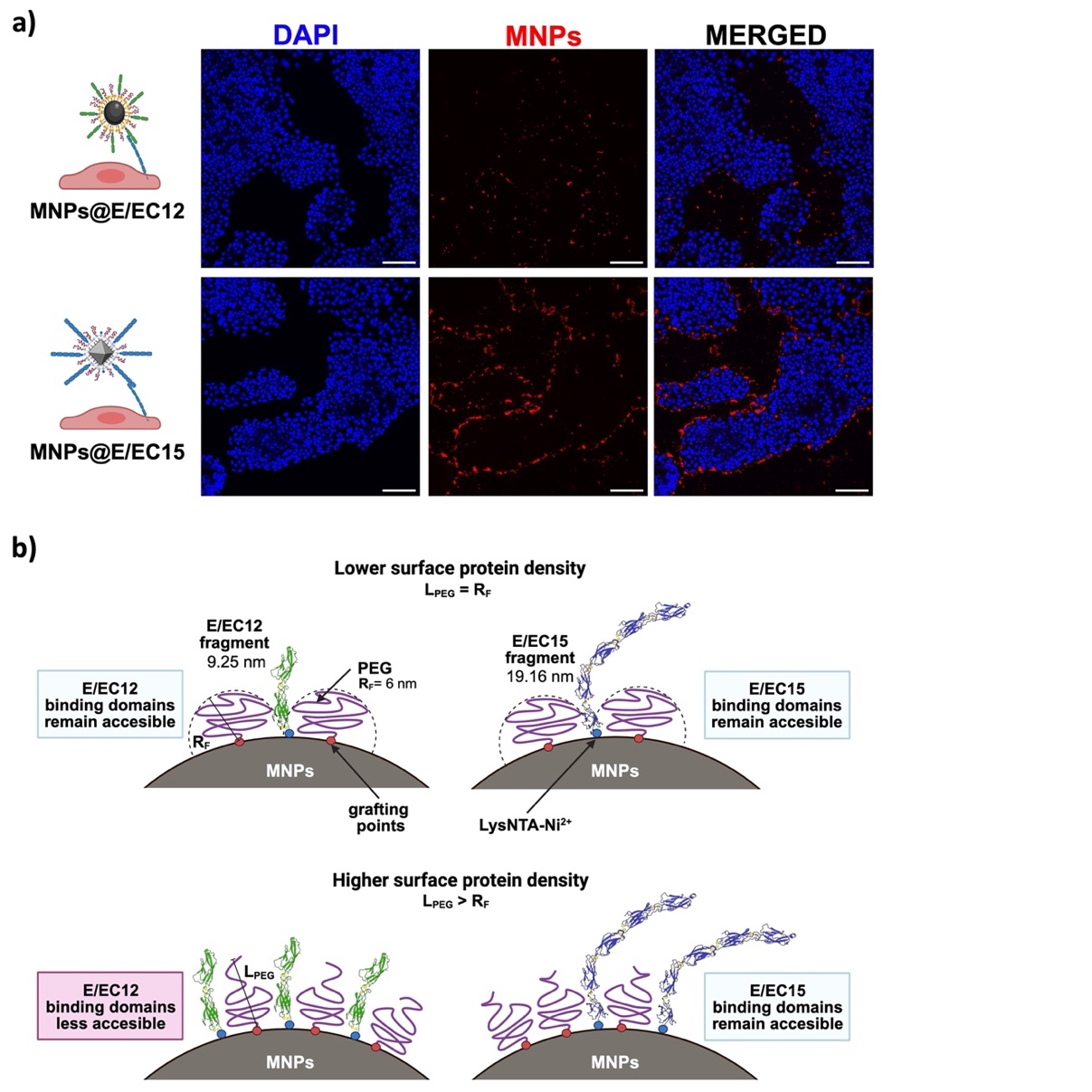

**Figure S3.** a) Labeling of MDCK cells with bioconjugates bearing E/EC12 (60 fragments/MNP) or E/EC15 (17 fragments/MNP) fragments. The bioconjugates containing E/EC12 fragments were obtained following the optimized conditions established in our prior report.^1^ Nuclei were stained with 4′,6-diamidino-2-phenylindole dilactate (DAPI; blue); MNPs containing TAMRA are shown in red; scale bar: 50 μm. b) Schematic representation of the proposed effect of surface protein density on E/EC12 or E/EC15 interaction. As the surface protein density increases, the length of PEG (L_PEG_) initially equal to the Flory radius (R_F_) also increases, limiting accessibility of the shorter E/EC12 fragments due to steric hindrance, thereby affecting bioconjugates interaction capabilities.

### 5. **Theoretical simulation of the magnetic field in the center of the cell culture dish during magnets movement**

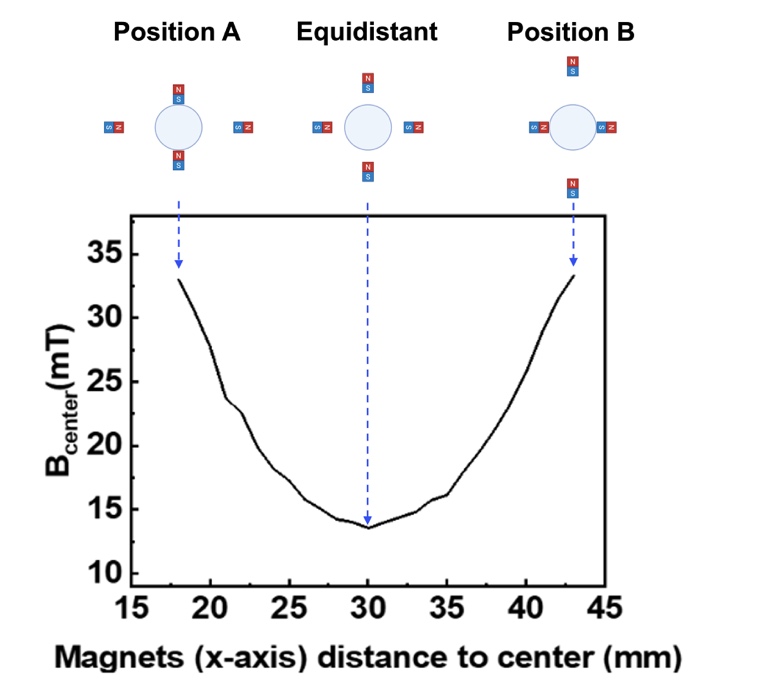

**Figure S4.** Theoretical calculation of the magnetic field intensity measured at the center of the petri dish during the full motion of the magnets from position A to position B. The maximum intensity is obtained when two sets of magnets are located closer to the petri dish (position A or B).

### 6. **Immunofluorescence of MDCK cells labeled with E-cadherin bioconjugates and stimulated with magnets**

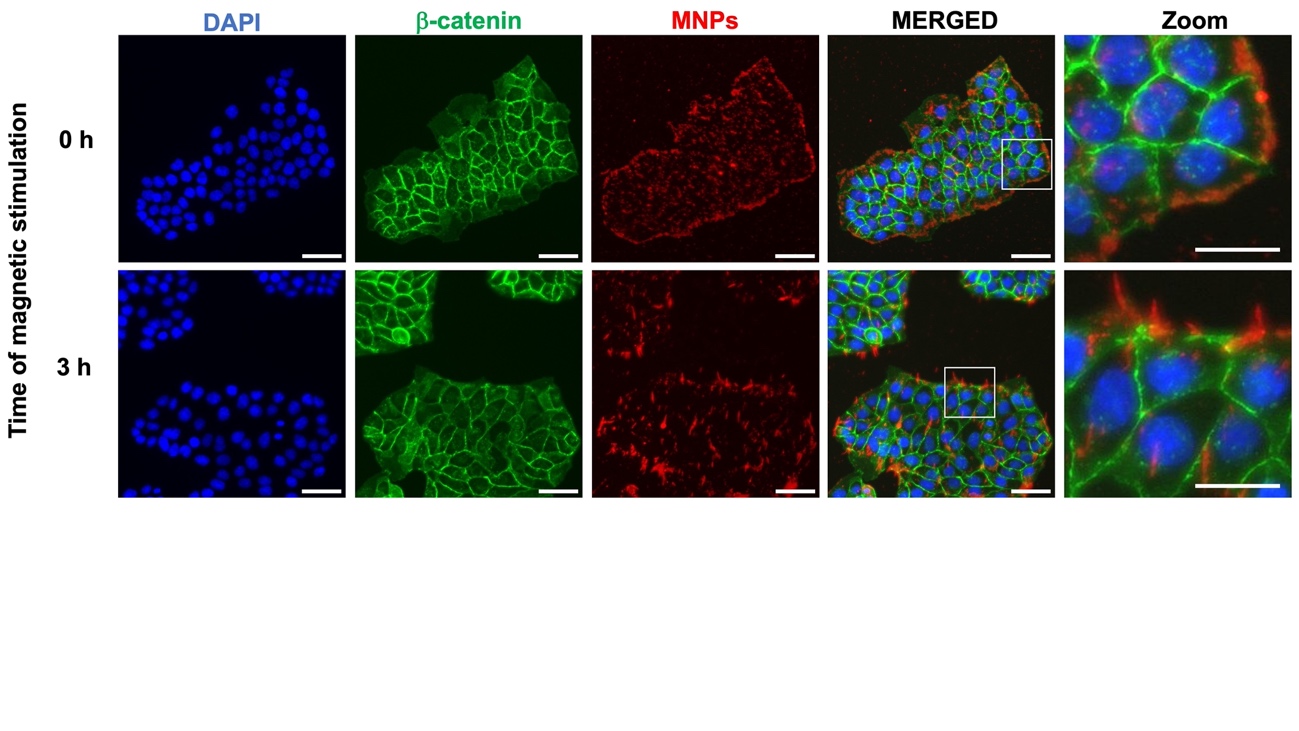

**Figure S5.** Fluorescence microscopy images of MDCK cells incubated 30 minutes with the bioconjugates, washed (0 h) and then stimulated with the magnets for three hours before fixation. Nuclei were stained with 4′,6-diamidino-2-phenylindole dilactate (DAPI; blue); β-catenin was immunostained and is shown in green (AF488); MNPs containing TAMRA are shown in red; scale bar: 50 µm; Zoom scale bar: 25 µm.

### 7. **RNA-seq intra- and intergroup variability and DEGs**

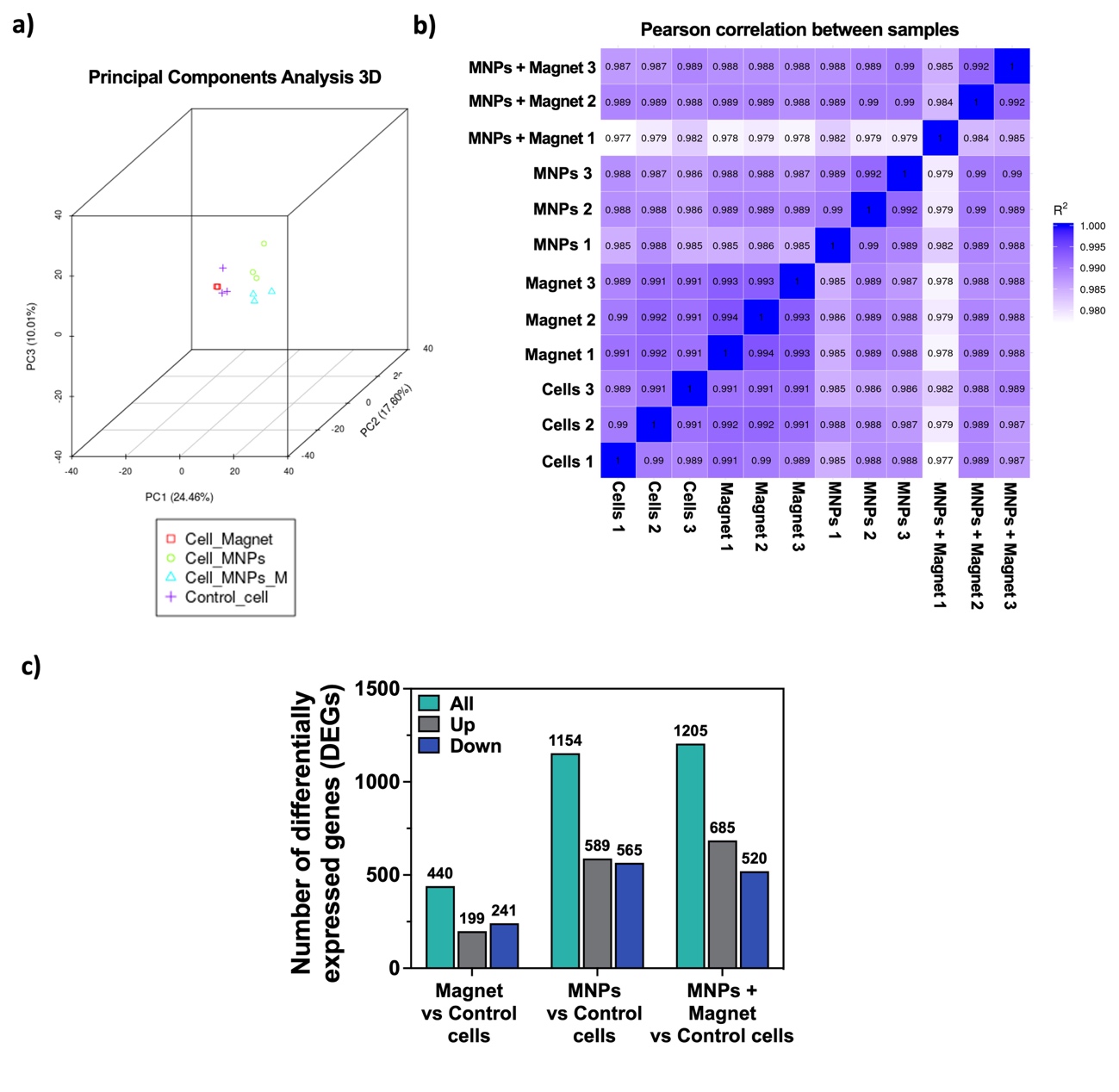

**Figure S6.** a) Principal components analysis (PCA) in three dimensions, showing intra- and intergroup variability. b) Pearson correlation coefficients (PPC) indicating intra- and intergroup variability. Lower intragroup variability (higher PPC), corresponding to lower variability among replicates, and higher intergroup variability (lower PPC) were observed between the four evaluated treatments. c) Number of DEGs resulting from the treatment versus control comparisons.

### 8. **Functional analysis: MNPs vs Control comparison**

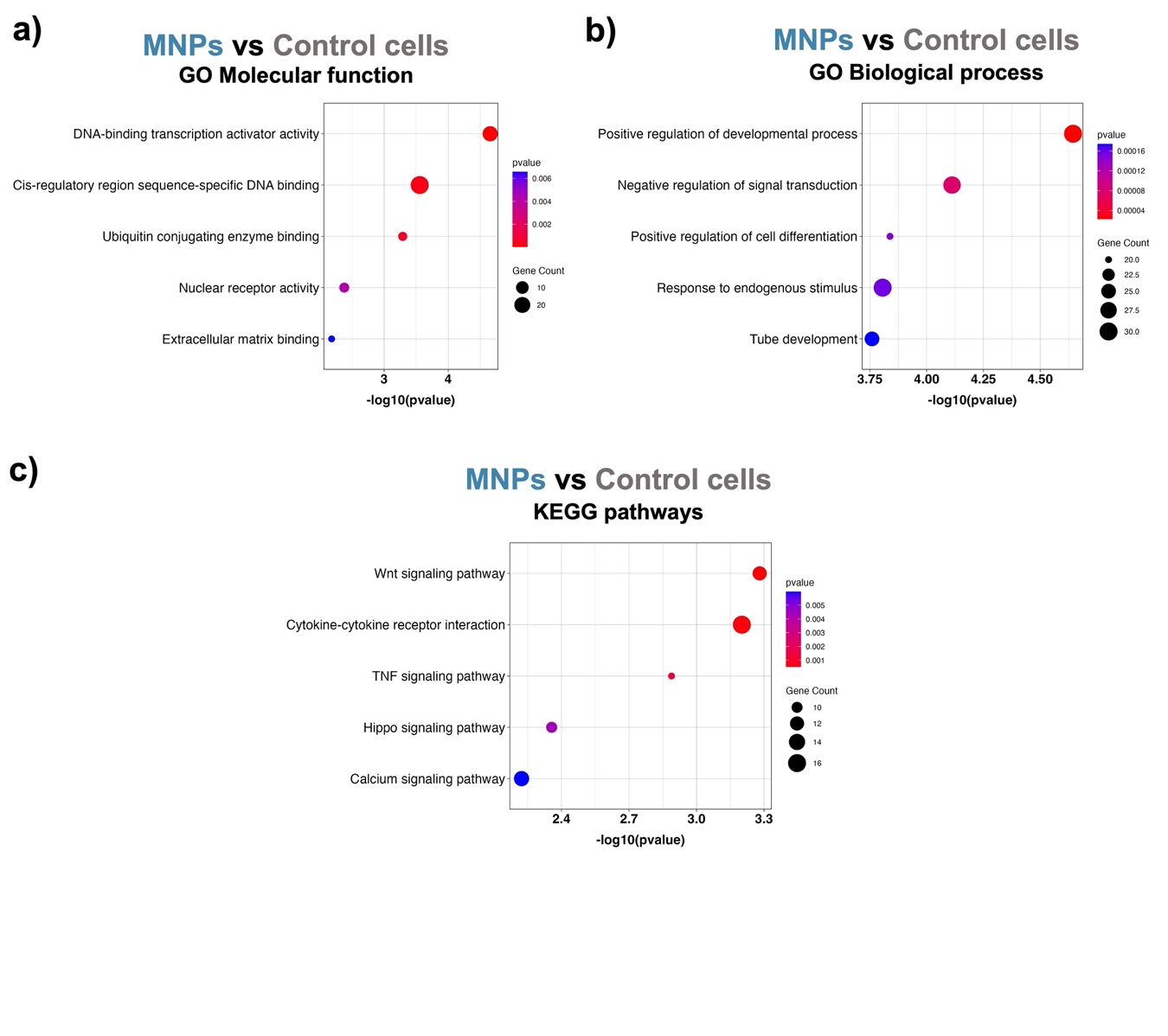

**Figure S7.** GO enrichment analysis of the 647 DEGs identified from MNPs vs Control comparison. a) GO analysis showing the top 5 molecular functions (MF) enriched by the presence of MNPs. b) GO analysis showing the top 5 biological processes (BP) enriched by the presence of MNPs. c) KEGG pathway enrichment analysis showing the top 5 pathways enriched.

### 9. **Functional analysis: MNPs + Magnet vs Control comparison**

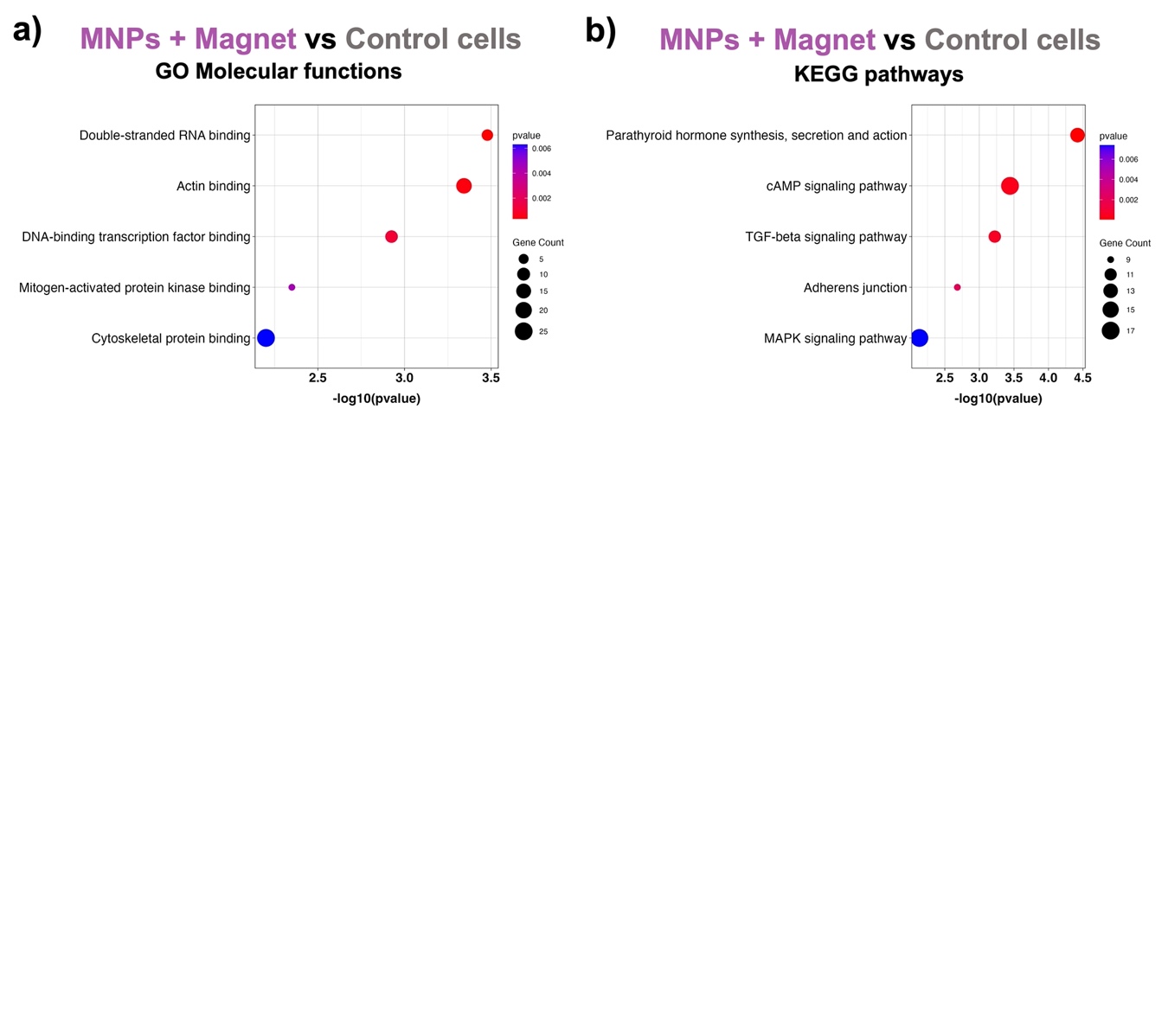

**Figure S8.** GO enrichment analysis of the 704 DEGs identified from MNPs + Magnet vs Control comparison. a) GO analysis showing the top 5 molecular functions (MF) enriched by the presence of MNPs combined with magnetic stimulation. b) KEGG pathway enrichment analysis showing the top 5 pathways enriched.

### 10. **‘MNPs + Magnet’ vs MNPs comparison**

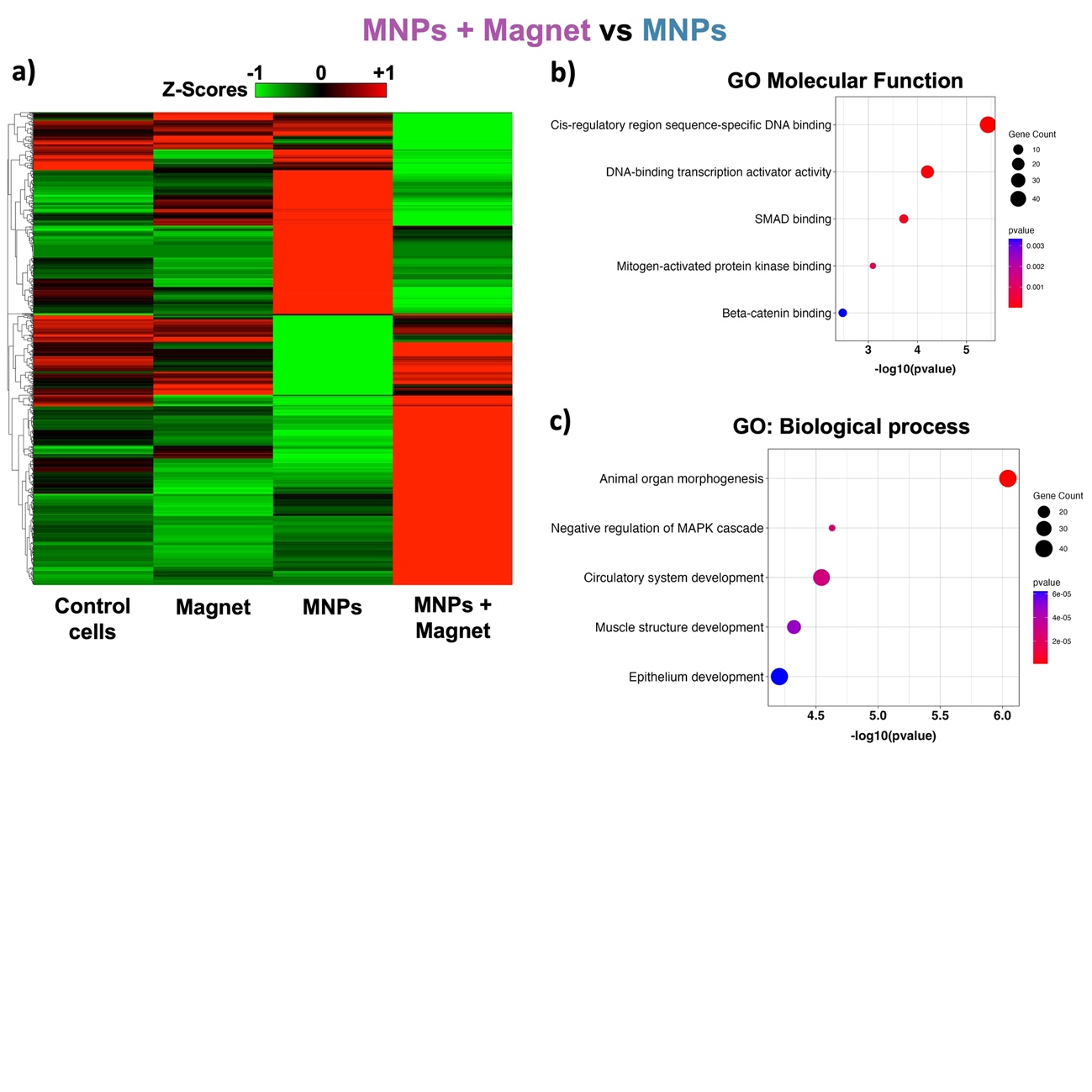

**Figure S9.** Analysis of the 1004 DEGs found in ‘MNPs + Magnet’ vs MNPs comparison. a) Heatmap of the 1004 DEGs identified. b) and c) GO enrichment analysis: Biological process and Molecular function categories respectively, showing the top 5 enriched terms resultant.

### 11. **RT-qPCR**

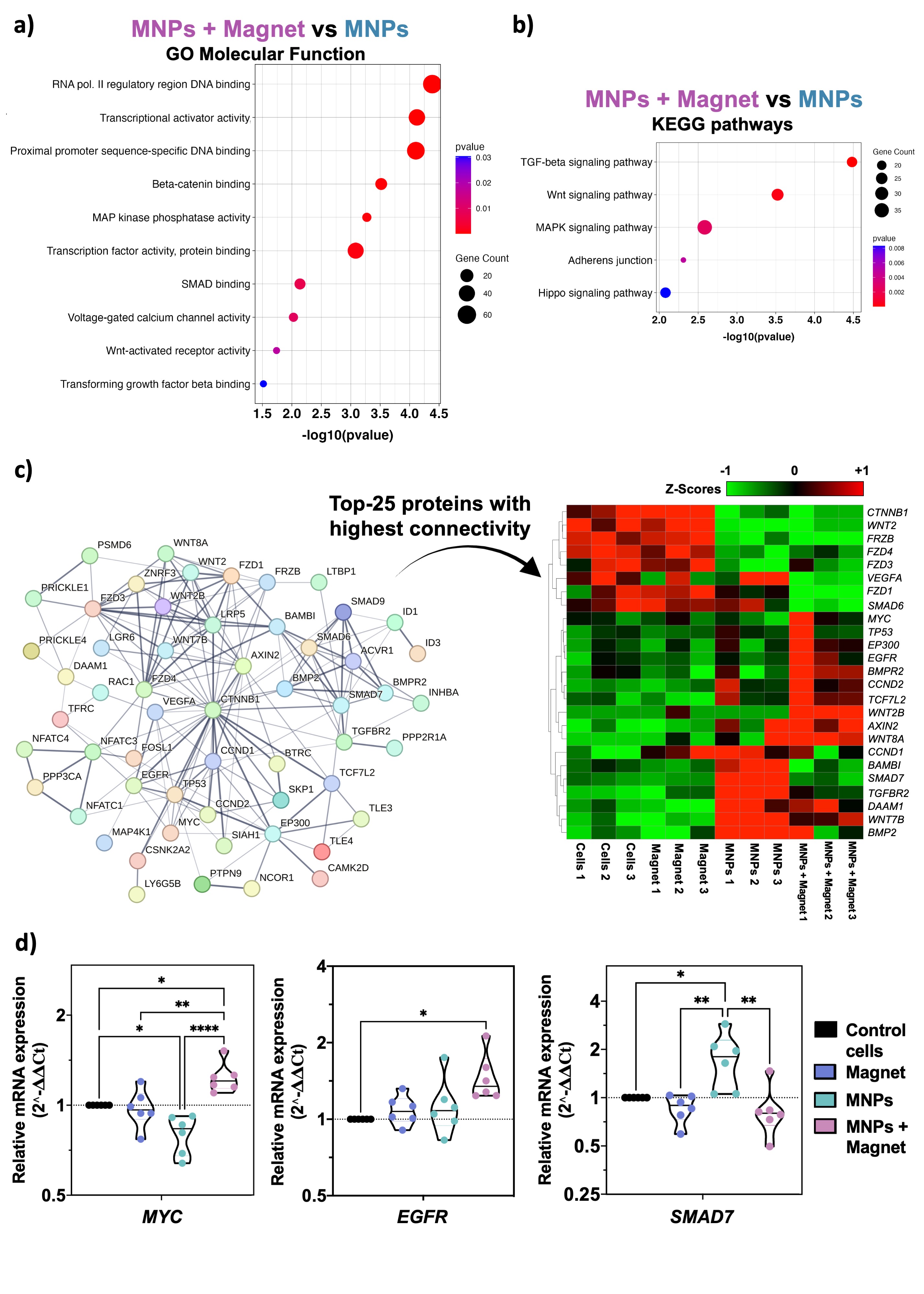

**Figure S10.** Real-time quantitative polymerase chain reaction (RT-qPCR) analysis showing relative mRNA expression levels (2^-ΔΔCt) of *MYC*, *EGFR* and *SMAD7* for validation of RNA-seq results (N=6 independent experiments performed in different days). Black asterisks indicate statistical differences (*p < 0.05; **p < 0.01; ***p < 0.001; ****p < 0.0001). One-way ANOVA followed by Tukey’s multiple comparison test were performed.

12. **Flow cytometry analysis of E-cadherin expression**

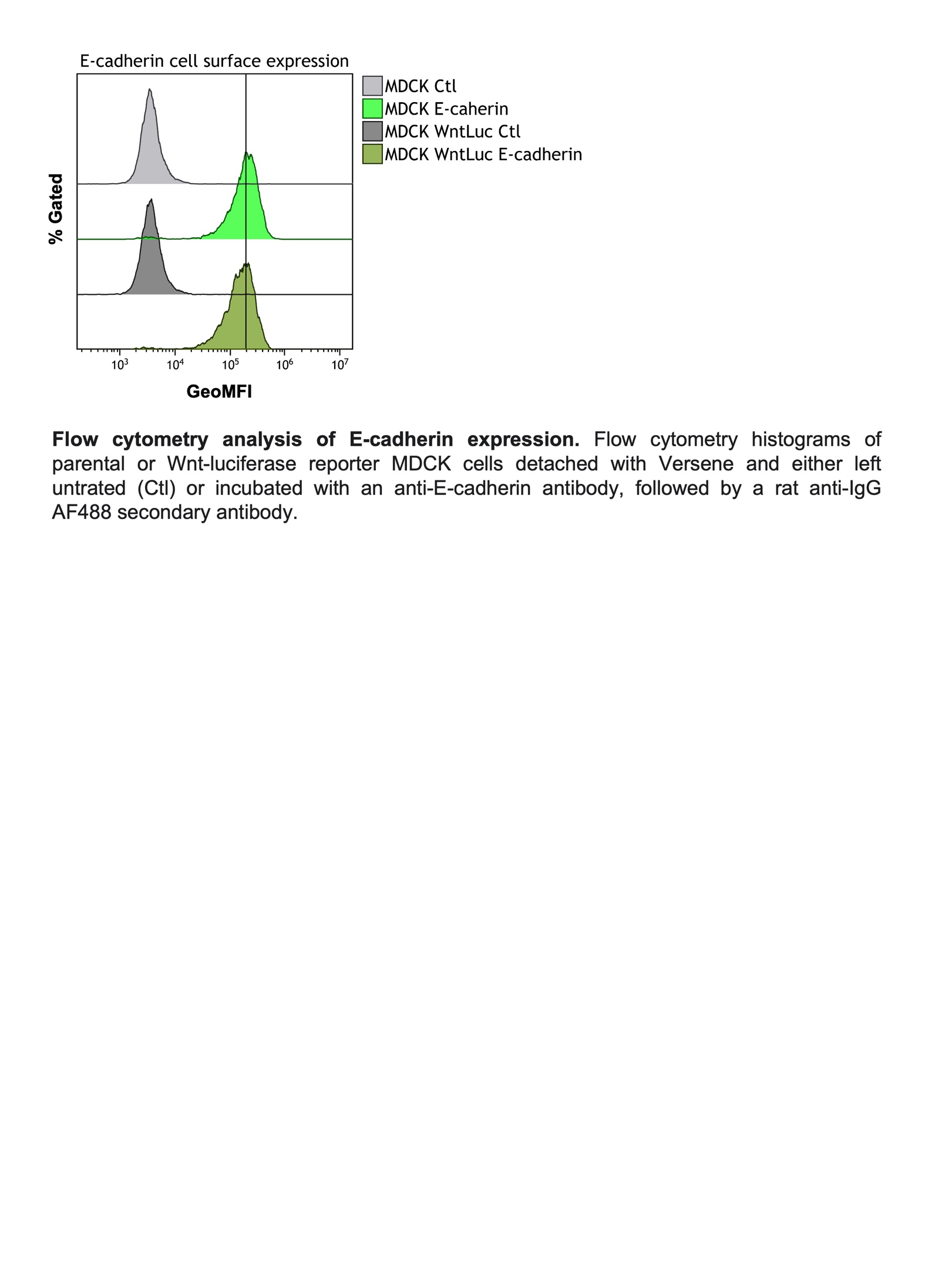

**Figure S11.** Flow cytometry histograms of parental or Wnt-luciferase reporter MDCK cells detached with Versene and either left untreated (Ctl) or incubated with an anti-E-cadherin antibody, followed by a rat anti-IgG AF488 secondary antibody. GeoMFI: Geometric mean fluorescence intensity.

### 13. **EdU staining**

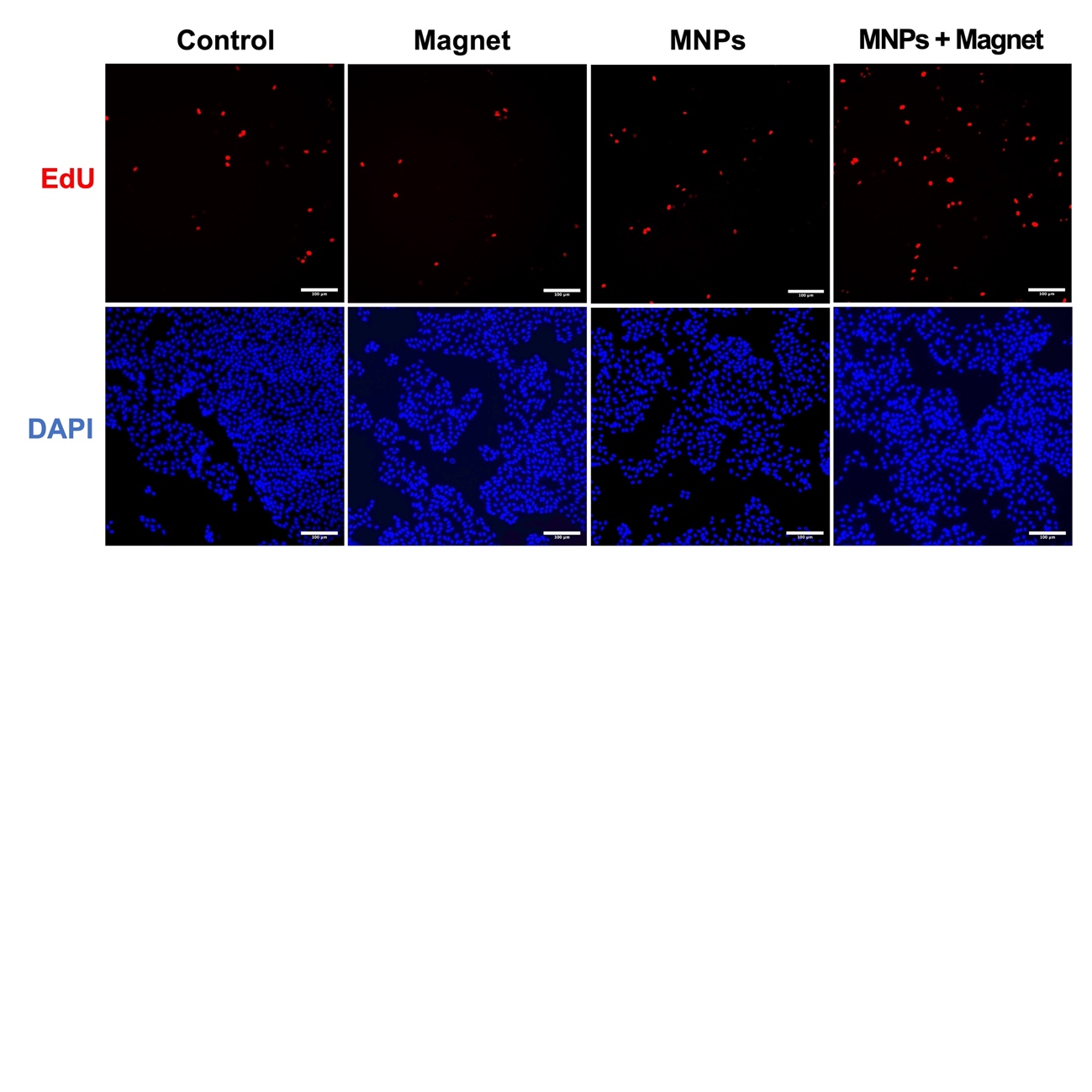

**Figure S12.** MDCK cells after EdU staining. EdU-positive cells are shown in red; nuclei are shown in blue. Scale bar: 100 μm.

### 14. **Gap closure curve**

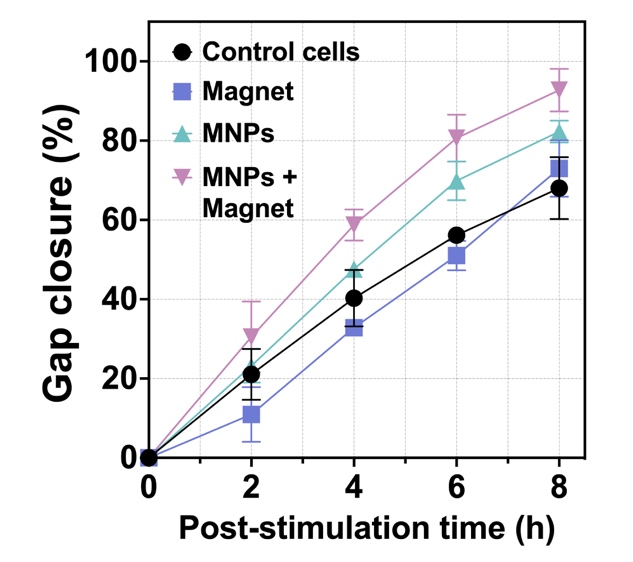

**Figure S13.** Kinetics of gap closure. The slope from 0-6 hours post-stimulation was used to calculate the corresponding cell front velocity (μm min^-1^).

### 15. **Supplementary materials and methods**

**RT-qPCR**

Total RNA was extracted following the same method described in the manuscript for RNA sequencing. The sequences of the primers corresponding to the *MYC*, *EGFR* and *SMAD7* genes are provided in Table S2. Primers were purchased in the lyophilized format and reconstituted in PCR-grade water following manufacturer’s instructions. RNA concentration and purity were analyzed on a Biotek Synergy H1 UV/vis microplate spectrophotometer measuring absorbance at 260, 280 and 320 nm, and by agarose gel electrophoresis (1.5 % m v-1) stained with Gel Red in 0.5x tris borate EDTA (TBE) buffer. The cDNA was synthesized from 1 μg of RNA using a high-capacity cDNA reverse transcription kit following manufacturer’s instructions.

RT-qPCR was carried out using NZYSupreme qPCR Green Master Mix in a CFX-OPUS 96 Real Time PCR System (BioRad). The following conditions were used: polymerase activation at 95 ºC for 2 min; initial denaturation at 95 ºC for 30 seconds; 40 extension cycles (5 seconds at 95 ºC and 25 seconds at 60 ºC); melting curve (to evaluate the formation of unwanted amplification products): 0.5 ºC increments from 60 to 95 ºC in 5 seconds steps. The expression of the different genes was evaluated according to the 2^-ΔΔCt method, using GAPDH gene as housekeeping gene. Each RT-qPCR reaction was performed in triplicate, and six independent experiments were evaluated.

**Table S2.** Primer sequences used for RT-qPCR of mRNA extracted from MDCK cells

| **Gen** | **Forward primer**  **5' to 3'** | **Reverse primer**  **5' to 3'** | **Amplicon size** | **Reference** |
| --- | --- | --- | --- | --- |
| ***EGFR*** | CCACCTGCGTGAAGAAATGC | CTTACACTTGCGGACACCATC | 116 | 2 |
| ***MYC*** | TCGCCTATTTGGGAAGACAC | AAGCTGACGTTGAGAGGCAT | 141 | 3 |
| ***SMAD7*** | GGACAGCTCAACTCGGACAA | ATGGAGAAACCGGGGAACAC | 203 | Designed de novo |
| ***GAPDH*** | GTCCCCACCCCCAATGTATC | TCCGATGCCTGCTTCACTAC | 98 | 4 |

### 16. **Calculations**

**S1 calculation: Wound healing assay**

The gap closure percentage was determined using the following equation:

$Gap closure \%=\left( \frac{A_{t=0}-A_{t=\Delta t}}{A_{t=0}} \right)*100\%$.

The front velocity (μm min^-1^) was calculated with the following equation:

$$V=\left( \frac{Rate of gap closure}{Lenght of cell front* Number of migrating cell fronts} \right)x100\%$$

The rate of gap closure (μm^2^ min^-1^) was obtained calculating the slope of a trend line obtained from image S10 with the data from 0 to 6 hours post-stimulation. The length of cell front is the height or length of the image, in our case 1430 $\mu m$.

### 17. **References**

(1) Castro-Hinojosa, C.; Del Sol-Fernández, S.; Moreno-Antolín, E.; Martín-Gracia, B.; Ovejero, J. G.; de la Fuente, J. M.; Grazú, V.; Fratila, R. M.; Moros, M. A Simple and Versatile Strategy for Oriented Immobilization of His-Tagged Proteins on Magnetic Nanoparticles. Bioconjug. Chem, 34 (12), 2275–2292, (2023). https://doi.org/10.1021/acs.bioconjchem.3c00417.

(2) Zorzan, E., et al. Whole-Transcriptome Profiling of Canine and Human in Vitro Models Exposed to a G-Quadruplex Binding Small Molecule. Sci Rep 8, 17107 (2018). https://doi.org/10.1038/s41598-018-35516-y.

(3) Angstadt, A., et al. Characterization of canine osteosarcoma by array comparative genomic hybridization and RT-qPCR: Signatures of genomic imbalance in canine osteosarcoma parallel the human counterpart. Genes Chromosomes & Cancer Vol. 50, Issue 11, 859-875 (2011). https://doi.org/10.1002/gcc.20908.

(4) Carvajal-Agudelo, J., et al. Evaluation of gamma-actin, beta-actin, GAPDH, and 18S as reference genes for qRT-PCR using blood samples in canine mammary research. Veterinarska stanica, Vol. 53 No. 1, (2022). <https://doi.org/10.46419/vs.53.1.5>.
